# Metabolic reprogramming underlies cavefish muscular endurance despite loss of muscle mass and contractility

**DOI:** 10.1101/2022.03.12.484091

**Authors:** Luke Olsen, Michaella Levy, J Kyle Medley, Huzaifa Hassan, Richard Alexander, Emma Wilcock, Kexi Yi, Laurence Florens, Sean McKinney, Robert Peuß, Jenna Persons, Alexander Kenzior, Ernesto Maldonado, Andrew Gluesenkamp, Edward Mager, David Coughlin, Nicolas Rohner

## Abstract

Physical inactivity – specifically the lack of moderate-to-vigorous activity – is a scourge to human health, promoting metabolic disease and muscle wasting. Interestingly, multiple ecological niches have relaxed investment into physical activity, providing unique evolutionary insight into adaptive physical inactivity. The Mexican cavefish *Astyanax mexicanus* lost moderate-to-vigorous activity following cave colonization, reaching basal swim speeds ~3-fold slower than their river-dwelling counterpart. We found that this was accompanied by a marked shift in body composition, decreasing muscle mass by 30% and increasing fat mass by 40%. This shift persisted at the single muscle fiber level via increased lipid and sugar accumulation at the expense of myofibrillar volume. Transcriptomic analysis of laboratory-reared and wild-caught cavefish indicated this shift in investment is driven by increased expression of *pparγ* – the master regulator of adipogenesis – with a simultaneous decrease in fast myosin heavy chain expression. *Ex vivo* and *in vivo* analysis confirmed these investment strategies come with a functional trade-off, decreasing cavefish muscle fiber shortening velocity, time to maximal force, and ultimately maximal swimming velocity. Despite this, cavefish displayed a striking degree of muscular endurance, reaching maximal swim speeds ~3.5-fold faster than their basal swim speeds. Multi-omics analysis indicated metabolic reprogramming, specifically increased phosphoglucomutase-1 abundance, phosphorylation, and activity, as contributing mechanisms enhancing cavefish glycogen utilization under metabolically strenuous conditions. Collectively, we reveal broad skeletal muscle reprogramming following cave colonization, displaying an adaptive skeletal muscle phenotype reminiscent to mammalian disuse and high-fat models while simultaneously maintaining a unique capacity for sustained muscle contraction under fatiguing conditions.

**Significance:** The evolutionary consequence of decreased physical activity upon skeletal muscle physiology remains unexplored. Using the Mexican cavefish, we find loss of moderate-to-vigorous swimming following cave colonization has resulted in broad shifts in skeletal muscle investment – away from muscle mass and instead toward fat and sugar accumulation – ultimately decreasing muscle fiber twitch kinetics. Surprisingly though, cavefish possessed marked muscular endurance, reaching maximal swimming speeds rivaling their river-dwelling counterpart. Multi-omics analysis revealed carbohydrate metabolic reprogramming as a contributing component, most notably elevated abundance and phosphorylation of the glycogenolytic enzyme Phosphoglucomutase-1 – a likely adaptation to cave-specific hypoxia. These findings emphasize the impact multiple selective pressures have on skeletal muscle physiology, providing the first evolutionary insight into skeletal muscle adaptation following decreased activity.

## Introduction

Throughout evolution, movement and feeding have shaped skeletal muscle physiology – whether by enhancing calcium transients to power “super-fast” contraction in the Toadfish swimbladder, accumulating massive swaths of mitochondria to permit large-scale migration in the Atlantic Bluefin Tuna, or restructuring joint anatomy to power explosive snapping in the Pistol Shrimp (Rome et al. 1996; Dumesic et al. 2019; Kaji et al. 2018). Indeed, extreme environmental conditions necessitate exaggerated skeletal muscle phenotypes, providing powerful insight into the boundaries of skeletal muscle adaptation and performance. While insightful, research into these “locomotor extremes” (Dickinson et al. 2000) have remained at a single end of the locomotory spectrum. In fact, in contrast to the examples above, many species have drastically decreased physical activity levels, at times remaining completely inactive for months and possibly even years (Fröbert et al. 2020; Balázs et al. 2020; Hendrickson et al. 2001). Troglobites are prime examples of such extreme changes to locomotion, with multiple cave-dwellers remaining either motionless, or adapting a glide-and-rest swimming behavior, in stark contrast to their above-ground counterparts (Hüppop et al. 2000). While adaptive, these extreme feats of ‘laziness’ beg the question of how skeletal muscle responds to prolonged periods of inactivity and may ultimately shed light on the evolutionary consequence of physical inactivity on skeletal muscle physiology - a current scourge to human health (Booth et al. 2017). To this point, we established the Mexican cavefish as a comparative model to provide an evolutionary perspective into skeletal muscle physiology following dramatic changes in locomotion.

The tetra species *Astyanax mexicanus* is found throughout Mexico and South Texas and is comprised of river-dwelling surface fish and cave-dwelling cavefish. Approximately 160,000 years ago, ancestral surface fish invaded surrounding caves, resulting in multiple independently evolved cave populations (Gross, 2012; Herman et al. 2018). Many of these colonized caves are completely devoid of light, resulting in diminished biodiversity and food availability.

Consequently, cavefish have adapted a suite of morphological, metabolic, and behavioral traits permitting survival in their nutrient-depleted conditions. For example, cavefish have evolved metabolic strategies such as hyperphagia, enhanced fat storage, insulin resistance, and hyperglycemia (Xiong et al. 2018; Riddle et al. 2018; Aspiras et al. 2015). Additionally, because these cave communities typically have only a single stygobitic vertebrate species (cavefish), they no longer face predation by other species. As a result, cavefish have dramatically changed their swimming behavior, abandoning energetically expensive burst-like swimming and instead relying on slow, continuous movement (Elipot et al. 2013; Carlson et al. 2018). Thus, the Mexican cavefish have adapted a suite of metabolic and behavioral traits similar to human metabolic disease, providing unique evolutionary insight into the consequence of such phenotypes on skeletal muscle physiology (Rohner, 2018). Importantly, cave and surface populations are conspecific and can be raised under identical, controlled laboratory conditions, permitting comparative studies of heritable physiological consequences to decreased strenuous movement following cave colonization. As such, here we leverage the *Astyanax mexicanus* system to address the consequence of, and adaptation to, distinct environmental conditions on skeletal muscle physiology following cave colonization – most notably changes to physical activity – providing unique evolutionary insights into the extremes of skeletal muscle adaptation.

## Results and Discussion

### Shift in swimming speed and body composition following cave colonization

Cavefish have repeatedly lost aggressive and territorial behavior comprised of burst-like swimming, and instead rely on slow, continuous movement (Elipot et al. 2013; Carlson et al. 2018). While previous studies have investigated the average velocities between surface fish and multiple cavefish populations (Carlson et al. 2018), quantitative analysis of maximal swimming speeds during unperturbed tank swimming – an often-challenging task due to the aggressive behavior of surface fish – have not been performed. To address this, recordings of laboratory-reared surface fish (originating from the Rio Choy) and laboratory-reared cavefish (originating from two independently colonized caves – Pachón cave and Tinaja cave) (Fig. 1A) were manually cropped at times containing burst-like swimming (fastest swimming bouts in cavefish), and frame-by-frame distances were analyzed (see methods, Movies S1A-C, and Data S1). This experimental design revealed both Pachón and Tinaja cavefish swim markedly slower during their fastest swimming bouts, reaching average burst velocities (acceleration-to-deceleration) ~3-fold slower (Fig. 1B) and maximal velocities (greatest frame-to-frame distance covered) ~2-fold slower (Fig. S1A) than surface fish. These findings are consistent with Paz et al. (2020), who found cavefish have reduced angular velocity during a C-start (escape) response, differences observed as early as 6 days post fertilization. Providing support that these differences are not a consequence of laboratory conditions, video recordings of wild cavefish (Pachón cave) and wild surface fish (Rio Choy) swimming revealed surface fish incorporate sustained moderate swimming – a form of rheotaxis required to remain stationary against the river current – interspersed by vigorous burst-like swimming (Movie S2A). In stark contrast, cavefish displayed an uninterrupted glide-and-rest swimming behavior (Movie S2B), confirming cave colonization resulted in a behavioral phenotype lacking both moderate and vigorous swimming.

**Figure 1.**
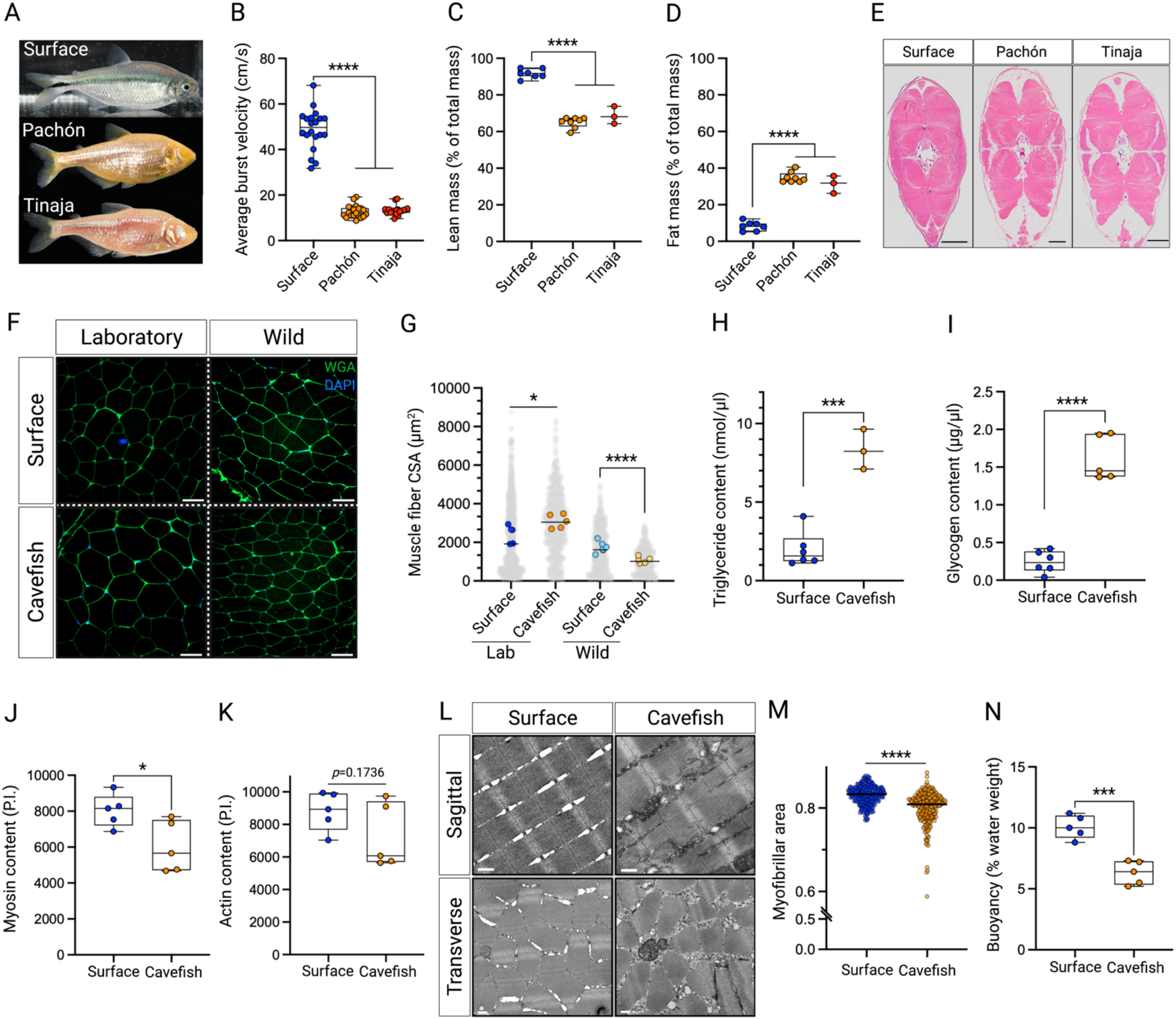
Shift in cavefish swimming speed and body composition. (A) Images of surface fish and two independently evolved cavefish populations: Pachón and Tinaja. (B) Average burst velocity of surface fish, Pachón, and Tinaja cavefish (n=20, 20, and 18, measurements per population, respectively). Percent (C) lean mass and (D) fat mass of *A. mexicanus* using an echoMRI (n=8 for Pachón and surface, n=3 for Tinaja). (E) Representative full body transverse cross-sections of *A. mexicanus* used for the echoMRI measurements showing skeletal muscle (pink) and subcutaneous fat (white) (scale bar = 500 μm). (F) Muscle fiber cross-sections from laboratory-reared (left) and wild-caught (right) surface fish and cavefish (Pachón). Muscle fibers are demarcated via wheat germ agglutin (green) (scale bar = 50μm). (G) Muscle fiber cross-sectional area (CSA) of laboratory-reared and wild-caught surface fish and cavefish (Pachón). Light grey circles represent individual muscle fibers. Colored circles represent mean muscle fiber cross-sectional area for each sample (n=5 for lab Pachón, lab surface, and wild Pachón, n=6 for wild surface). (H) Triglyceride content (nmol/μl) (n=3 and 6 for Pachón and surface, respectively). (I) Glycogen content (μg/μl) (n=5 and 6 for Pachón and surface, respectively). (J) Myosin content in pixel intensity (P.I.) (n=5 per population). (K) Actin content in pixel intensity (P.I.) (n=5 per population). (L) Transverse and sagittal electron micrographs of surface fish and cavefish (Pachón) skeletal muscle (n=2 per population, scale bar = 500nm). (M) Quantification of myofibril area. Each point indicates the relative area of myofibrils within a single EM image. ~150 images were quantified per sample. See methods for further detail. (N) Percent water weight of surface fish and cavefish (Pachón). The higher the percent denotes greater weight in water. Significance for Fig. 1B-D were calculated with a one-way ANOVA with Bonferroni FDR correction. For all other analysis, significance was calculated with unpaired student’s t-test. Data is presented as ±SEM, \**p*<0.05, \*\*\**p*<0.001. \*\*\*\**p*<0.0001.

Skeletal muscle contraction is a key stimulus influencing muscle physiology (Franchi et al. 2017), with multiple disuse models revealing increased fat accumulation and decreased skeletal muscle mass, fiber size, and force production following physical inactivity (Rudrappa et al. 2016). Because cavefish have markedly reduced their swimming speed following cave colonization (Fig. 1B and S1A), we hypothesized that their skeletal muscle would reflect an “inactive” phenotype, and hence shift investment away from muscle mass and instead toward fat accumulation. To this point, baseline *A. mexicanus* body composition was gathered using an echo magnetic resonance imaging (echoMRI) system (Xiong et al. 2021). Supporting our hypothesis, we found both Pachón and Tinaja cavefish have ~2.5-fold less lean mass (Fig. 1C) and ~4-fold more fat mass (Fig. 1D) than surface fish. These findings were confirmed via histological analysis of full body cross-sections (Fig. 1E), revealing a near identical shift away from cavefish muscle mass and instead toward fat accumulation (for quantification see Fig. S1B and S1C). To our surprise, the decrease in cavefish muscle mass occurred independent of a reduction in muscle fiber size – a predominant factor contributing to muscle atrophy following physical inactivity (Bodine. 2013). In fact, we found Pachón cavefish had ~22% larger muscle fiber cross-sectional area than surface fish (Fig. 1F and 1G). Curious whether laboratory conditions – specifically regular feeding and constrained movement – drives this difference between surface fish and cavefish muscle fiber size, we collected wild *A. mexicanus* from the Pachón cave (cavefish) and Rio Choy (surface fish) and measured their muscle fiber cross-sectional area. Intriguingly, the opposite phenotype was observed - specifically, wild cavefish had ~30% smaller muscle fibers than wild surface fish, and ~66% smaller muscle fibers than laboratory cavefish (Fig. 1F and 1G). While muscle fibers in wild surface fish tended to be smaller than laboratory surface fish, the degree of change was dramatically smaller when compared to cavefish populations (23% vs. 66%, respectively). These data indicate that, while cavefish have reduced investment into relative muscle mass (Fig. 1C and 1E), their muscle fibers possess a surprising degree of flexibility, most notably the capacity to hypertrophy, dependent upon environmental conditions - surpassing the flexibility of surface fish.

We next sought to explore the underlying mechanism(s) contributing to this unique degree of muscle fiber hypertrophy within cavefish. In brief, muscle fiber hypertrophy takes place through two primary mechanisms: i) sarcoplasmic hypertrophy - the increase in muscle fiber size via increasing the non-contractile sarcoplasmic volume (i.e. fats, sugars, and sarcoplasmic proteins) and ii) myofibrillar hypertrophy - the increase in muscle fiber size via myofibrillar accretion (i.e. sarcomeric proteins actin and myosin) (Roberts et al. 2020). Because cavefish can store considerable energy reserves throughout their body (Xiong et al. 2021), we reasoned their muscle fiber growth arises through sarcoplasmic hypertrophy at the expense of myofibrillar hypertrophy - specifically via accumulation of sugars and fats. Supporting our hypothesis, we found cavefish skeletal muscle had ~4-fold greater triglyceride levels (Fig. 1H) and ~7-fold greater glycogen levels (Fig. 1I) than surface fish. Intriguingly, analysis of a recent lipidomic and metabolomic dataset of *A. mexicanus* skeletal muscle revealed cavefish preferentially accumulate saturated fatty acids, ceramides, and sphingoid bases, while simultaneously decreasing investment in free amino acids, most notably the mTORC1-activating leucine - a lipid and amino acid profile linked to muscle atrophy and diminished contractility (Rivas et al. 2019; Medley et al. 2020; Fig. S1D). Reflecting this, fractionation experiments found that cavefish skeletal muscle have an ~25% and ~17% (*p*=0.1736) decrease in the contractile protein’s myosin and actin, respectively (Fig. 1J and 1K). Electron micrographs supported these findings (Fig. 1L), revealing cavefish muscle fibers have significantly fewer myofibrils with a subsequent increase in sarcoplasmic space relative to surface fish (Fig 1M).

Notably, we noticed that cavefish muscle fibers have distinct aggregates of glycogen granules both within the intermyofibrillar space (between myofibrils) and intramyofibrillar space (between contractile filaments), with very few granules identified within surface fish (Fig. 1L). These findings are particularly interesting since teleost muscle glycogen stores are presumed to reflect their activity levels – such as the highly active tuna possessing 8-fold more glycogen than the sedentary carp (Sänger et al. 2001) – suggesting cavefish face alternative stimuli resulting in muscle fiber glycogen accumulation (discussed in following sections). Intriguingly, we found the unique cavefish body composition coincided with an elevated water buoyancy (i.e. weigh less in water) relative to surface fish (Fig. 1N). We reason this is a consequence of their decreased investment in dense tissue (myofibrils) and increased investment in buoyant tissue (fats). In fact, while the cavefish dry weight was ~20% greater than surface fish (6.08g vs 4.84g, respectively), their water weight was ~22% less than surface fish (0.38g vs. 0.49g, respectively), suggesting a thrifty skeletal muscle remodeling strategy incorporated by cavefish to both increase energy reserve capacity while simultaneously improve swimming economy via enhanced static lift.

### Change in cavefish body composition is reflected at the level of gene expression

Having established a clear shift in cavefish skeletal muscle composition following cave colonization, we next sought to determine whether this change was reflected at the level of gene expression. To this end, we conducted RNA-sequencing of surface fish and cavefish skeletal muscle (FDR<0.01) followed by Gene Ontology (GO) enrichment analysis of the differentially expressed genes (DEG’s) (Fig. 2A-C and Data S2). This revealed several overexpressed pathways related to muscle contractility and structural integrity - specifically “myosin complex” and “actin cytoskeleton” (Fig. 2C). Interestingly, most genes within these pathways were downregulated within cavefish, most notably genes encoding fast myosin heavy chain proteins - a class of myosin heavy chains with the capacity for rapid sarcomere cross-bridge cycling (Schiaffino et al. 2011). In fact, of the DEG’s within the “myosin complex” pathway, ~70% were downregulated within cavefish, decreasing between 6-to-50 fold - all of which were fast myosin heavy chain genes (Fig. S2A, Data S3). In addition, DEG’s contributing to muscle atrophy (*gadd45ga* - Fontes-Oliveira et al. 2013) and swimming speed via dopamine inhibition (*mblac1* - Hardaway et al. 2015) were increased and decreased, respectively, within cavefish (Fig. S2B and S2C). Confirming this transcriptome signature is a cave-specific phenomenon, we sequenced the skeletal muscle from an additional, independently evolved cavefish population (Tinaja cavefish), and found a similar decrease in fast myosin heavy chain expression (Fig. 2D and Data S4), along with increased and decreased *gadd45ga* and *mblac1* expression, respectively (Fig. S2B and S2C), indicating a conserved decrease in expression of genes necessary for rapid muscle contraction following cave colonization

**Figure 2.**
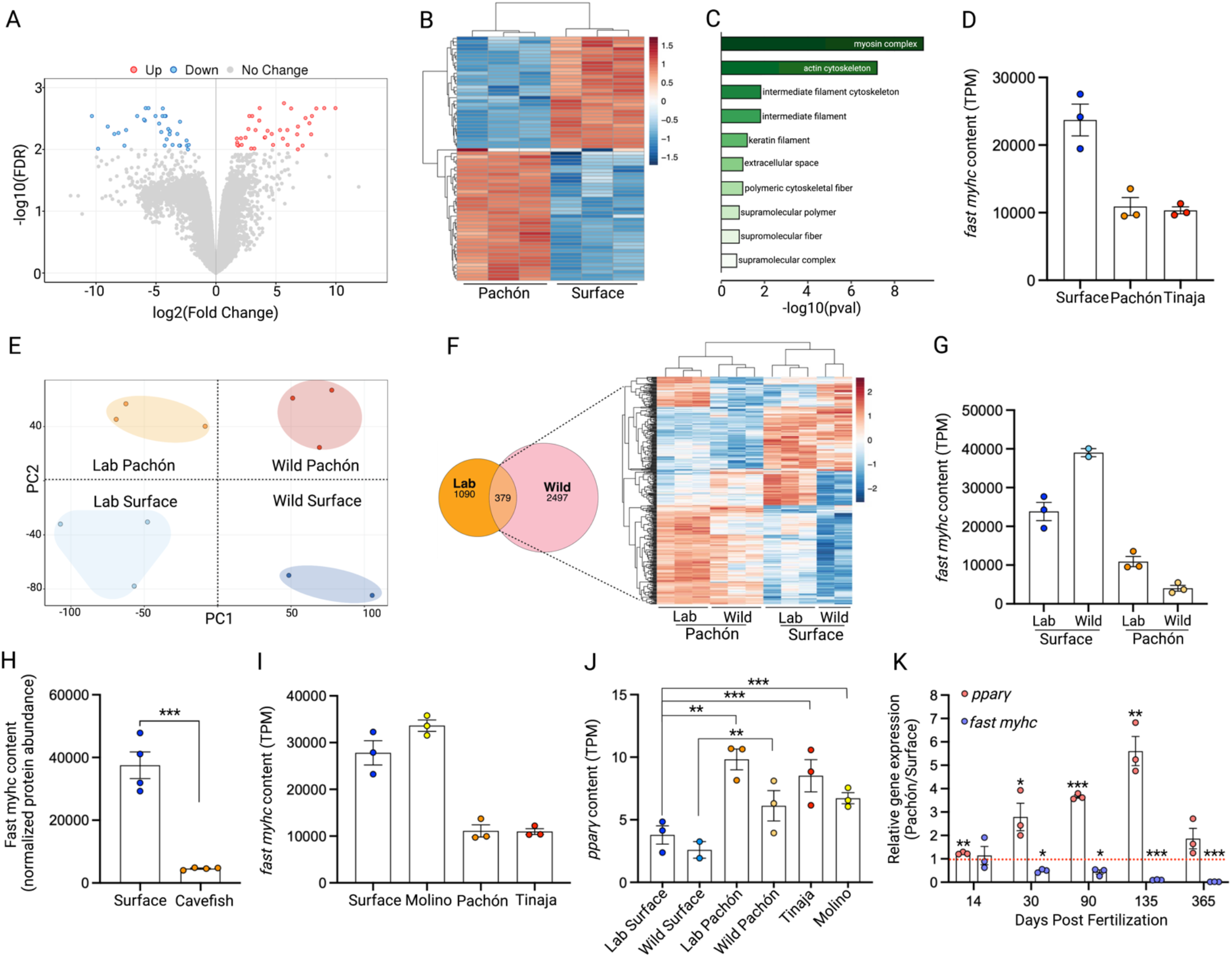
The cavefish skeletal muscle transcriptome reflects their body composition. (A) Volcano plot of the differentially expressed genes (DEG’s) between laboratory-reared surface fish and cavefish (Pachón) (up- and down-regulated in Pachón fish relative to surface fish). (B) Heatmap of the differentially expressed genes showing individual replicate data. (C) Gene Ontology enrichment analysis of the DEG’s from Fig. 2A and Fig. 2B. (D) Cumulative abundance of shared differentially expressed fast myosin heavy chain genes in Tinaja and Pachón relative to surface fish (TPM: transcripts per million). (E) Principal component analysis of the laboratory-reared and wild-caught Pachón and surface fish transcriptome. (F) Venn diagram of all DEG’s between laboratory-reared and wild-caught fish. Specific emphasis is placed on the 379 overlapping genes with their expression shown in the adjacent heatmap. (G) Cumulative abundance of the shared differentially expressed fast myosin heavy chain genes between laboratory-reared and wild-caught Pachón and surface fish. (H) Cumulative abundance of the differentially expressed fast myosin heavy chain proteins between cavefish (Pachón) and surface fish (n=4 per population). (I) Cumulative abundance of fast myosin heavy chain genes of those identified as significantly different between laboratory-reared Pachón vs laboratory-reared surface fish. These genes (Data S6) were then used to determine cumulative fast myosin heavy chain abundance in Molino and Tinaja. (J) *pparγ* expression between all fish populations. (K) *pparγ* and fast myosin heavy chain (*fast-myhc*) expression across developmental timepoints. Expression is taken relative to surface fish (indicated by the red line). Statistical analysis for RNA-sequencing is described in the methods. For Fig. 2K, data was analyzed via unpaired students t-test for each timepoint. Data is presented as ±SEM. \**p*<0.05, ** *p* < 0.01, \*\*\**p*<0.001.

To determine whether the above findings are a consequence of domestication, specifically constrained movement due to laboratory housing, we collected additional wild *A. mexicanus* from the Pachón cave and wild surface fish from the San Antonio River and conducted RNA-sequencing of their skeletal muscle. Principal component analysis (PC) revealed a clear discrimination between both environmental conditions (wild/laboratory – PC1) and fish ecotypes (Pachón/surface fish – PC2) (Fig. 2E), consistent with previous comparative transcriptome datasets (Krishnan et al. 2020). Using a relaxed FDR threshold of <5% to broadly detect changes in gene expression, we compared the DEG’s of the wild samples (2876 DEG’s) against the DEG’s of the laboratory samples (1469 DEG’s), revealing 379 overlapping DEG’s between populations (Fig. 2F and Data S5) – the majority differentially expressed in the same direction. GO-term analysis of these 379 genes again revealed “actin cytoskeleton” and “myosin complex” as the top ranked differentially regulated pathways (Fig. S2D). Similar to our laboratory transcriptome dataset, 80% of the genes within the “myosin complex” pathway were downregulated within wild cavefish – decreasing from 2.6-to-85 fold – all of which were fast myosin heavy chain genes (Fig. 2G and Data S5). Validating our RNA-seq data, global proteomic analysis of whole muscle lysate from cavefish and surface fish found an ~8-fold decrease in fast myosin heavy chain protein abundance in cavefish compared to surface fish (Fig. 2H), denoting a conserved reduction at both the transcript and protein level.

The above findings are crucial to account for the impact environmental factors have on skeletal muscle gene expression and confirm that the skeletal muscle phenotypes observed within laboratory cavefish are largely representative of cavefish in the field. However, we questioned whether these phenotypes might reflect phenotypic plasticity and not genetic inheritance. Specifically, might the decreased swimming speed of cavefish in both laboratory and wild conditions be the sole contributor to their reduced “contractile” skeletal muscle phenotype – a possible reflection of the “use it or lose it” phenomenon (Wisdom et al. 2015). To address this, we conducted RNA-sequencing of a phylogenetically younger cavefish population originating from the Molino cave (termed Molino cavefish). Molino cavefish are considered a “new” *A. mexicanus* cavefish lineage – having colonized their cave ~110,000 years ago – and genomically cluster more closely to surface fish than to Pachón or Tinaja cavefish (Herman et al. 2018). Importantly though, Molino cavefish have converged on similar swimming behaviors to Pachón and Tinaja cavefish, lacking territorial and aggressive behavior and thus lacking vigorous swimming (Elipot et al.

2013). With this unique mix of genetic and behavioral traits, we reasoned that if the decrease in fast myosin heavy chain expression is solely a result of phenotypic plasticity (i.e. slow swimming), fast myosin heavy chain expression within Molino cavefish should decrease regardless of their genetic background. Intriguingly, in contrast to the phenotypic plasticity hypothesis, Molino cavefish possessed ~3-fold greater fast myosin heavy chain levels than both Pachón and Tinaja cavefish - levels similar to surface fish (Fig. 2I). These findings indicate that the decreased fast myosin heavy chain expression in Pachón and Tinaja is likely not a result of phenotypic plasticity but instead is driven by genetic inheritance. However, it should be noted that a subset of lowly expressed fast myosin heavy chain isoforms were shifted in a cave-like direction within Molino (i.e. shifted in a similar direction to Pachón and Tinaja), reflecting minimal, but not absent, myosin heavy chain divergence following Molino cave colonization (Data S6).

### Cavefish show increased expression of genes involved in skeletal muscle lipid metabolism

In contrast to decreased fast myosin heavy chain expression, cavefish skeletal muscle showed a consistent, and significant, 2.3- to 3.2-fold increase of the master regulator of adipogenesis *pparγ*. This was observed in both wild and laboratory cavefish - including Pachón, Tinaja, and Molino - relative to wild and laboratory surface fish (Fig. 2J). In fact, a timecourse qPCR analysis found cavefish gradually increase skeletal muscle *pparγ* expression as early as 14 days post fertilization which continued to adulthood, in contrast to the temporal dynamics of fast myosin heavy chain expression which declined early in development and persisted to adulthood (Fig. 2K). These findings support our recent discovery of Tinaja, Pachón, and Molino cavefish encoding a truncated variant of Period 2 - affecting its inhibitory *pparγ*-binding domain - a variant we validated is present in cavefish skeletal muscle (Xiong et al. 2021). Congruently, cavefish muscle had elevated expression of the lipid metabolism genes adiponectin b (*adipoqb*), ceramide synthase 4-b (*cers4b*), adiponutrin (*pnpla3*), and leptin receptor (*lepr*) (Fig. S2E-H). Many of these genes are adipocyte-specific, indicating cavefish skeletal muscle possess elevated intramuscular adipocytes, a phenomenon associated with muscle fiber atrophy and muscle weakness (Biltz et al. 2020). Collectively, these data support a cavefish skeletal muscle profile indicative of a shift away from skeletal muscle mass/contractility and instead toward fat accumulation, findings which strongly reflect their body composition in Figure 1.

### Functional analysis of *A. mexicanus* skeletal muscle

The skeletal muscle phenotype described above has repeatedly shown to result in muscle fiber functional decline – most notably diminished muscle fiber contraction – in both rodents and humans (Biltz et al. 2020; Choi et al. 2016). While this phenotype is likely beneficial within the starvation-prone cavefish, intramyocullar fat accumulation is poised to result in a functional decline in muscle fiber twitch kinetics. Seeking to address whether a similar physiological consequence occurs within cavefish, we conducted *ex vivo* muscle bundle contractility experiments. In brief, hypaxial myotomal muscle was dissected between the pectoral and pelvic girdle, and immediately placed in physiological saline. Isolated live muscle bundles (approximately 20 muscle fibers) were tied into a muscle mechanics chamber with a servomotor at one end and force transducer at the other, and platinum electrodes along each side of the chamber (Fig. 3A and Movie S3). This *ex vivo* approach is essential to exclude external confounding factors such as differences in neural innervation. Cavefish indeed demonstrated a trade-off in muscle contractility, having an ~18% reduction in muscle bundle shortening velocity (Fig. 3B), taking ~15% longer to reach maximal force (time from stimulus until maximum isometric force – Fig. 3C), and taking ~25% longer to relax following muscle contraction (time from maximum force to 50% relaxation – Fig. 3D) relative to surface fish.

**Figure 3.**
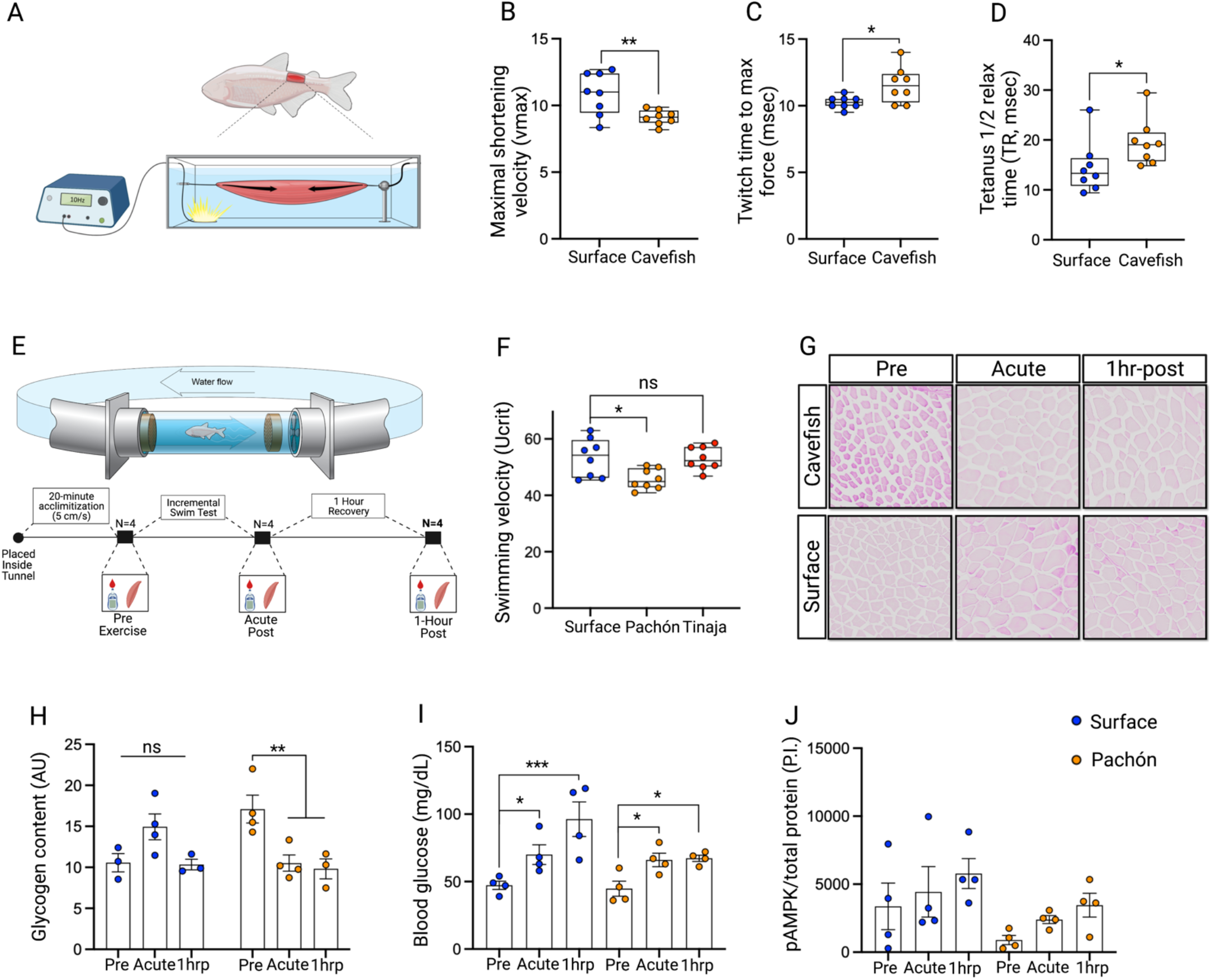
Cavefish maintain the capacity to increase swim speeds despite reduced muscle contractility. (A) Schematic of the muscle mechanics chamber showing site of skeletal muscle dissection, attachment, and stimulation. Arrows denote muscle shortening. (B) Maximal shortening velocity (vmax). (C) Twitch time to maximal force (msec). (D) Tetanus ½ relax time (TR, msec) (n=8 per population for each experiment. (E) Schematic of the swim tunnel with the experimental timeline of the incremental swim test and tissue/blood collection. (F) Maximal swimming velocity reached during the incremental swim test (Ucrit in cm/s) in surface fish and cavefish (Pachón and Tinaja) (n=8 per population). (G) Histological images of muscle fiber glycogen via Periodic acid-Schiff stain at the pre, acute-post, and 1-hour post timepoint of surface fish and cavefish (Pachón). (H) Quantification of glycogen content shown in Fig. 3G (n=3-4 per timepoint). (I) Blood glucose levels of exercised and non-exercised surface fish and cavefish (Pachón) (n=4 per timepoint). (J) Quantification of phosphorylated AMPK-Thr^172^ in the exercised and non-exercised surface fish and cavefish (Pachón) (n=4 per timepoint). Significance was determined with an unpaired students t-test (Fig. 3B-D), one-way ANOVA with Bonferroni FDR correction (Fig. 3F), and an ordinary, two-way ANOVA with Benjamini and Hochberg FDR correction (Fig. 3H-J). Data is presented as ±SEM, \**p* ≤ 0.05, \*\**p* ≤ 0.01, \*\*\**p* ≤ 0.001, ns = not significant).

### Cavefish maintain the capacity to increase swimming speeds under stimulated conditions

To test if the decreased *ex vivo* muscle contractility in cavefish results in decreased swimming velocities *in vivo*, we implemented an incremental swim test wherein cavefish and surface fish gradually (5 cm/s every 5 minutes) increase swimming speeds until volitional fatigue (Fig. 3E; Movies S4A and S4B). We found that cavefish reach maximal swimming velocities ~15% slower than surface fish (cavefish: 45.76 cm/s, surface fish: 53.41 cm/s – Fig. 3F). Surprisingly though, while slower than surface fish, cavefish showed an impressive ~3.5-fold capacity to increase their swimming speed between their incremental swim test (45.76 cm/s – Fig. 3F) and their average burst speed during tank swimming (12.84 cm/s - Fig. 1B), whereas surface fish remained largely unchanged between their incremental swim test (53.41 cm/s - Fig. 3F) and their average burst speed during tank swimming (48.86 cm/s - Fig. 1B). Differences can be seen in supplemental Figure 3A. Indeed, conducting the incremental swim test with an additional cavefish population (Tinaja cavefish) revealed a similar ~3.9-fold increase in maximal swimming velocity relative to baseline swimming speeds. In fact, Tinaja cavefish reached maximal swim speeds similar to surface fish (Tinaja cavefish: 53.11 cm/s), with no significant difference between populations (Fig. 3F, Fig. S3A, and Movie S4C). These data indicate cavefish have a marked ability to not only increase their swim speed under stimulated conditions but also maintain elevated swim speeds for extended periods of time (each swim test lasted ~40 minutes), a particularly striking display of muscular endurance for an animal irregularly exposed to such stimuli (discussed in more detail in the discussion).

### Cavefish have greater glycogen utilization during exercise

We next sought to characterize the molecular mechanisms underpinning cavefish muscular endurance. Because the incremental swim test served as an anaerobic stimulus, and since muscular endurance is a function of glycogen levels (Hermansen et al. 1967), we reasoned that the elevated glycogen stores within cavefish skeletal muscle (Fig. 1I) provide the needed metabolic substrate for sustained ATP production and muscle contraction, and hence render cavefish fatigue-resistant despite reduced muscle mass and contractility. To address this, we collected skeletal muscle from cavefish and surface fish before, immediately following (acute-post), and 1-hour following (1hrp) the incremental swim test (Fig. 3E). This experimental design confirmed cavefish have greater glycogen levels in the rested state, however, these levels significantly decreased (denoting utilization) both immediately following and 1-hour following exercise with no change in surface fish (Fig. 3G and 3H) - findings consistent with Salin et al. (2010) who found elevated cavefish glycogen utilization following metabolic perturbations relative to surface fish. To test if the incremental swim test served as a sufficient metabolic stimulus in both fish populations, we analyzed blood glucose levels at all timepoints and found both surface fish and cavefish significantly increased blood glucose at the acute and 1hrp timepoints (Fig. 3I). Additionally, we observed a similar, albeit insignificant, increase in phosphorylated AMPK-Thr^172^ - a common proxy of cellular energetic stress - in both surface fish and cavefish (Fig. 3J). Collectively, these data confirm cavefish utilized their elevated glycogen pool during the incremental swim test and, importantly, that this phenomenon is not a consequence of greater systemic metabolic perturbation in one fish population over the other. These findings thus suggest cavefish skeletal muscle has heightened intrinsic factors regulating both glycogen synthesis and degradation under divergent metabolic conditions.

### Multi-omics analyses suggest increased cavefish glycogen metabolism

To determine the cellular mechanisms underlying the enhanced cavefish glycogen metabolism, we performed unbiased proteomics via liquid chromatography-tandem mass spectrometry (LC/MSMS) on the exercised and non-exercised skeletal muscle samples described above (Fig. 4A and Data S7). Notably, conducting a KEGG pathway analysis of the top 50 ranked proteins identified “glycolysis/gluconeogenesis” as the most differentially regulated pathway with all proteins in this pathway increased in cavefish (Fig. 4B and Data S8). To confirm, and further expand upon our findings, we conducted a targeted proteomic analysis of the same samples and found half of the proteins within the glycolytic pathway (Aldoa, Eno1, Gapdh, Pfkm, Pgk1) were significantly increased within cavefish, along with the glycogen handling enzymes phosphoglucomutase 1 (Pgm1), glycogen phosphorylase (Pygm), and the enzyme lactate dehydrogenase (Ldh) (Fig. 4C and Data S9A) – findings similarly reflected in the increased expression of *pygm,pfkpb*, and *ldhba* in our transcriptomic dataset (Fig. S3B-D). Supporting these findings, analysis of a publicly available metabolomics dataset from *A. mexicanus* skeletal muscle (Medley et al. 2020) revealed a significant 1.7-fold increase of the glycolytic end-product pyruvate within cavefish skeletal muscle relative to surface fish (Fig. S3E). Taken together, these datasets confirm a cavefish skeletal muscle profile capable of elevated glycogen metabolism.

**Figure 4.**
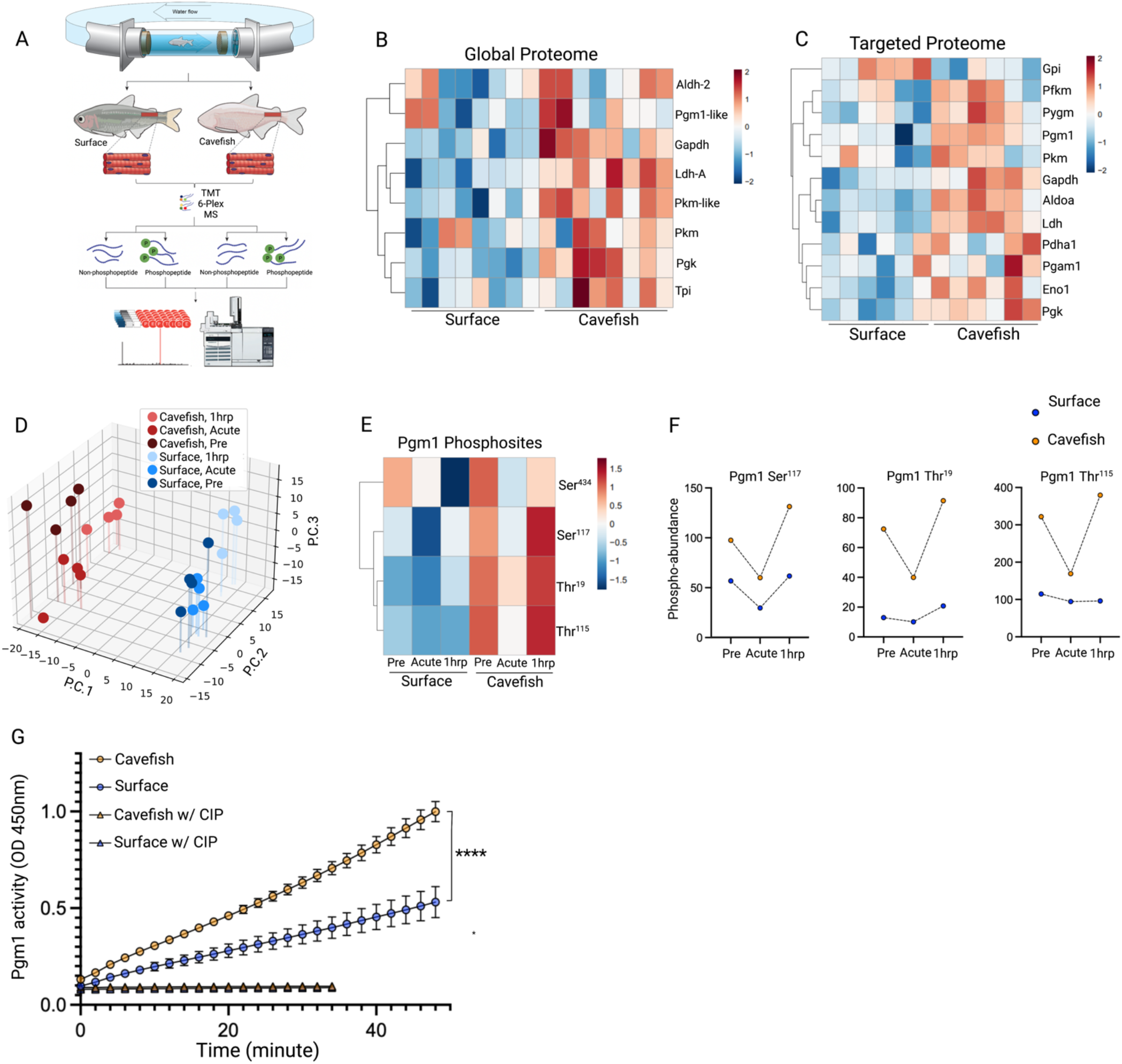
Multi-omics indicates increased abundance and activity of cavefish carbohydrate enzymes. (A) Tissue processing pipeline for both the proteomic and phosphoproteomic analysis for the surface fish and cavefish (Pachón) (n=12). (B) Heatmap from the global proteomic dataset of the proteins within the glycolysis/gluconeogenesis KEGG pathway (n=8 per population). (C) Heatmap of protein abundance levels from the targeted proteome analysis of proteins regulating carbohydrate metabolism. (D) Three-dimensional principal component analysis (P.C.) of all quantified phosphorylated peptides. (E) Heatmap of the mean peak intensity of Pgm1 phosphorylated sites at the pre, acute, and 1hrp timepoints. (F) Mean peak intensity of phosphorylation at each timepoint of surface fish and cavefish (Pachón) for Pgm1-Ser^117^, Pgm1-Thr^19^, and Pgm1-Thr^115^. (G) Pgm1 activity assay showing change in fluorescence over time in treated (with CIP) and untreated (without CIP) skeletal muscle samples from surface fish and cavefish (Pachón) (n=6 per population). Significance was determined using a repeated measures two-way ANOVA with Benjamini and Hochberg FDR correction (Fig. 4G). For proteomic data, analysis can be found in the methods. Data is presented as ±SEM, \*\*\*\**p* ≤ 0.0001.

### Cavefish skeletal muscle has increased Pgm1 phosphorylation and activity

While intriguing, we were hesitant to use static protein and metabolite levels as proxies for dynamic metabolic activity. Phosphorylation is a key regulator of metabolism and has been demonstrated to regulate carbohydrate metabolic dynamics (Chen et al. 2019), leading us to reason differential phosphorylation could more accurately reflect enzymatic flux, serving as an indirect proxy for glycogenolytic and glycolytic activity. To this point, a subset of the digested proteins used for proteomics were enriched for phosphorylated peptides via Sequential enrichment from Metal Oxide Affinity Chromatography (SMOAC) (Jae Choi et al. 2017) followed by LC/MSMS. This pipeline identified 430 phosphorylated peptides on 124 proteins in ≥2 biological replicates (Data S10). Three-dimensional principal component analysis (PC) of all phosphorylated peptides revealed a clear separation by population (PC1) and a less dramatic separation by timepoint (PC2; PC3) (Fig. 4D) - findings similarly reflected in human acute exercise omic datasets (Contrepois et al. 2020). With a particular interest in the elevated capacity for glycogen storage and utilization within cavefish, we focused on enzymes directly regulating glycogen metabolism. We considered Pgm1 a promising candidate since it i) acts in a bi-directional manner, capable of reversibly synthesizing or degrading glycogen dependent upon the energetic stimulus and ii) was identified as one of the most highly phosphorylated proteins at the pre timepoint in cavefish muscle relative to surface fish (Data S11). In fact, our phosphoproteomic data revealed that, of the four detected Pgm1 phosphorylated sites (Fig. 4E), three (Pgm1-Ser^117^, Pgm1-Thr^115^, Pgm1-Thr^19^) were increased at the pre timepoint 1.7-, 3-, and 5.5-fold, respectively. Phosphorylation levels on these sites were then uniformly decreased immediately following exercise (similar to recent findings in exercised rodents - Maier et al. 2021), followed by a sharp increase at the 1hrp timepoint - all of which were greater in the cavefish (Fig 4E and 4F). These findings indicate that in both basal and recovery states, cavefish have hyperphosphorylation of Pgm1 relative to surface fish - suggesting phosphorylation of Pgm1 is a crucial node in cavefish glycogen metabolism.

Previous work identified the above phosphorylation sites are essential for Pgm1 activity, with the phospho-serine Pgm1-Ser^117^ required for the inter-conversion between glucose-1-phosphate to glucose-6-phosphate (or vice-versa), thus serving as a pre-requisite for Pgm1 enzymatic activity (Lee et al. 2014). Notably, missense mutations (threonine-to-alanine) of Pgm1-Thr^115^ and Pgm1-Thr^19^ have been found in humans, resulting in exercise intolerance due to glycogenosis (Tegtmeyer et al. 2014; Stojkovic et al. 2009). As such, with elevated phosphorylation of all three phospho-sites in cavefish skeletal muscle, we reasoned they would in turn have increased Pgm1 activity. To test this, we measured Pgm1 activity of biopsied *A. mexicanus* skeletal muscle and found that cavefish have significantly greater Pgm1 activity relative to surface fish (Fig. 4G). Importantly, the enhanced cavefish Pgm1 activity was abolished following dephosphorylation via incubation with Calf Alkaline Intestinal Phosphatase (CIP) (Fig. 4G and S3F). In fact, we found the cavefish Pgm1 was more sensitive to dephosphorylation, with its activity reduced below surface fish levels following CIP incubation. These data strongly suggest phosphorylation is a crucial component driving the enhanced Pgm1 activity within cavefish skeletal muscle. Collectively, increases in both Pgm1 abundance (Fig. 4C) and phosphorylation levels (Fig. 4E and 4F) renders Pgm1 a likely candidate driving both increased cavefish glycogen content (Fig. 1I) and utilization (Fig. 3G and 3H) dependent upon the environmental conditions. Importantly, having elevated glycogen stores throughout the skeletal muscle – including within sarcomeres (Fig. 1L) – cavefish are poised to rapidly mobilize glycogen as a means to re-synthesize ATP and hence fuel prolonged muscle contraction under metabolically strenuous conditions.

The above data agree with van der Weele (2022) and Medley (2020) who found elevated carbohydrate transcripts and metabolites, respectively, within multiple cavefish tissues relative to surface fish. In fact, we recently found primary cavefish liver-derived cells have increased glycolytic capacity relative to surface fish liver-derived cells (Krishnan et al. 2022). We propose this is an adaptation to their hypoxic cave environment. Field studies from our lab have shown *A. mexicanus* caves have water oxygen saturation levels of 59% compared to 80% in surface fish waters (Rohner et al. 2013). Hypoxia is a strong selective pressure (Boggs et al. 2021; van der Weele et al. 2022), suggesting that this environmental stimulus has necessitated cavefish rely on anaerobic metabolism to supply the needed ATP for survival and, more specifically, increase investment in carbohydrate enzyme abundance and phosphorylation to meet their energetic demands. While further work is required to definitively address the effect differential phosphorylation has on *in vivo* glycolysis and ATP production, our proteomic and phosphoproteomic data strongly suggest a cavefish skeletal muscle profile well-adapted for carbohydrate metabolism - equipping cavefish with a surprising degree of muscular endurance under stimulated maximal swimming conditions despite loss of muscle mass and contractility - highlighting the diverse skeletal muscle adaptations following cave colonization.

## Conclusion

Our results highlight the remarkable breadths of skeletal muscle adaptation within the *Astyanax mexicanus*. Caves pose many challenges to survival - diminished food, decreased oxygen, and complete darkness - resulting in extreme behavioral, morphological, and physiological adaptations. Using surface and cave populations of *A. mexicanus*, we provide convincing evidence that a particularly important site of structural and metabolic reprogramming following cave colonization is the skeletal muscle. Cavefish have made large-scale shifts in their muscle investment, resembling disuse and high-fat conditions commonly seen within mammalian models (Booth et al. 2017). Indeed, with the loss of predators, water flow, and territorial behavior, cavefish have relaxed investment into skeletal muscle mass and contractility and instead shifted investment toward fat and sugar accumulation. Despite this, cavefish have adapted their metabolism to both increase and maintain elevated swimming speeds under stimulated conditions – findings we hypothesized result from elevated muscle glycogen levels and utilization during stimulated exercise. Follow-up proteomic and phosphoproteomic analyses of exercised cavefish skeletal muscle supported this hypothesis, revealing both protein and phosphorylation levels of key glycogenolytic and glycolytic enzymes increased within cavefish – a phenomenon we reasoned results from their hypoxic cave environment. In fact, we found hyperphosphorylation of the glycogenolytic/synthetic enzyme Pgm1 results in increased Pgm1 activity within cavefish skeletal muscle. This finding is particularly interesting since the degree of Pgm1 expression and phosphorylation strongly tracks with various metabolic diseases, most notably cancer and type 2 diabetes (Batista et al. 2020; Li et al. 2020). It is thus likely that chronic hyperphosphorylation of Pgm1 can be both physiological and pathological dependent upon the environmental conditions. Considering the hypoxic conditions under which cavefish evolved, Pgm1 hyperphosphorylation is likely adaptive, serving as a crucial node in cavefish skeletal muscle metabolism due to Pgm1 bi-directional properties – permitting both increased glycogen synthesis and breakdown dependent upon environmental conditions – providing a surprising degree of muscular endurance despite loss of skeletal muscle mass and contractility. However, in addition to hypoxia, periodic cave flooding, and thus elevated water flow, cannot be ruled out as a potential stimulus influencing cavefish skeletal muscle metabolism. In fact, occasional flooding within the Pachón and Tinaja cave can, at times, lead to swift water flow exiting the mouth of the cave. As such, adapting the ability to withstand periodic water flow to mitigate being swept away may serve as a crucial survival strategy.

## Acknowledgments

We would like to thank the cavefish facility at the Stowers Institute for cavefish husbandry support, specifically Zachary Zakibe, Molly Miller, Elizabeth Fritz, David Jewell, Franchesca Hutton-Lau, Andrew Ingalls, Diana Baumann, and Adam Petrie. We would like to thank Sylvie Rétaux and Mathilda Mommersteeg for providing wild *A. mexicanus* videos. We thank Mark Miller for assistance with illustrations. We thank all core support provided by the Stowers Institute; specifically, Seth Maloney, Dai Tsuchiya, and Nancy Thomas of Histology, Cindy Maddera of Microscopy, and Rhonda Egidy and Amanda Lawlor of Molecular Biology. We would like to especially thank all members of the Rohner Lab for the critical review of our manuscript along with Dr. Gustavo Blanco, Dr. Matt Morris, and Dr. Chad Slawson for their critical comments throughout the planning and experimental process. NR is supported by institutional funding, NIH Grant 1DP2OD028806-01, NIH Grant R01 GM127872, NSF IOS-1933428 and NSF EDGE award 1923372.

## Methods

### Laboratory fish husbandry

*Astyanax* husbandry and care was conducted identical to Xiong et al. (2021) and was approved by the Institutional Animal Care and Use Committee (IACUC) of the Stowers Institute for Medical Research on protocol 2021-122. All methods described here are approved on protocol 2021-129. Housing conditions meet federal regulations and are accredited by AAALAC International.

### Video tracking and tank burst swimming

Video tracking took place in the Pentair AES ZHAB fish rack Enclosure #LC60-C-A. 3-liter Pentair Aquatic Ecosystems #PCT3 tanks were modified to have internal acetal panels on front, bottom, and rear for improved video tracking. Current-USA TrueLumen Pro LED Aquarium Light Model 3031 was used for lighting. Videos were taken with a Logitech C920 webcam and exported to Open Broadcaster Software on a connected laptop. Three fish (either Pachón, Tinaja, or surface) were placed in a 3-liter water tank with continuous water circulation. Fish were placed inside the recording enclosure for 2 weeks prior to recording to acclimate to the surroundings. Fish were then recorded for 2 hours. Each video was marked at specific times containing fastest fish movements. For surface fish, this consisted of true bursts – aggressive behavior between two fish. For Pachón and Tinaja fish, the fastest swimming bouts often consisted of accelerations following a turn. Each swim clip began at the beginning of acceleration and concluded at the beginning of deceleration. ffmpeg was used to cut the videos into 4-second clips (2 seconds before and after the specified time for a total of 120 frames). Videos were then converted to flat image files (.tiff) and ROI’s were placed at the front edge of the fish of interest using FIJI/ImageJ. ROI files were calculated using Python analyzing the instantaneous velocity in each frame of the image. The scripts for breaking the MP4 videos into clips, and calculating the velocity from ROI.zip files can be found here: https://gist.github.com/richard-alexander/e3bded51a9e23dfccc1afc2c16fa58a2. Average burst velocity was measured at the beginning of an acceleratory burst and concluded at the onset of deceleration. Maximal burst velocity was the greatest distance covered between two frames.

### Wild sample collection and tissue processing

Wild fish collection in Mexico were collected as previously described (Krishnan et al. 2020) and conducted under permit No. SGPA/DGVS/03634/19 granted by the Secretaría de Medio Ambiente y Recursos Naturales to Ernesto Maldonado. Samples of wild surface fish were collected from the San Antonio River at the San Antonio Zoo, Texas. For RNA-sequencing, skeletal muscle was dissected immediately posterior to the dorsal fin and submerged in RNAlater until further analysis (1-2 weeks from initial collection). Immediately upon arrival to the final destination, skeletal muscle was gently blotted to remove RNAlater and placed in 1mL of Trizol (Ambion). Subsequent processing for RNA extraction and sequencing is identical to that described below (*Lab fish RNA-sequencing*). For histological analysis, skeletal muscle was dissected from the same region as that used for RNA-seq (separate fish) and immediately submerged in 100% ethanol. Tissue remained in ethanol until further processing (~2 weeks). Muscle fiber processing for cross-sectional area measurements was then conducted identical to that described below (*Lab fish muscle fiber size*).

### EchoMRI™

~1 year old fish were used to measure body composition using the EchoMRI™ analyzer. For female fish, eggs were removed prior to the assay. Technical replicates were measured and averaged as the readout for each sample. Total lean mass and total fat mass was normalized to total body weight and indicated as percent lean mass and percent fat mass.

### Histology processing for EchoMRI™ samples

Following processing with the EchoMRI™, fish were measured for length/weight and muscle was dissected across the entire truck – approximately 0.5cm from the base of the tail – and placed in 4% paraformaldehyde (PFA) at 4°C for 5 days. The tissue was subsequently washed in 1xPBS (pH 7.4) for 2×15 minutes at room temperature and then placed in 15% sucrose-1xPBS overnight. The samples were then moved to 30% sucrose-1xPBS at 4°C followed by hematoxylin & eosin staining. Samples were imaged with a VS120 virtual slide microscope (Olympus) and analyzed with ImageJ. To quantify muscle mass, the muscle area was taken relative to the total body area within each section. Care was taken to only use high quality sections avoiding any tissue damage. Two technical replicates were used for each biological replicate.

### Lab fish muscle fiber size

Skeletal muscle was collected from cavefish and surface fish similar in size. Muscle tissue was fixed in 4% PFA for 72hrs at 4°C followed by 2×6 minute washes in 1xPBS at room temperature and dehydrated by serial incubations in ethanol. Tissue samples were then embedded in paraffin and sectioned at a thickness of 10μm. Sections were deparaffinized, rehydrated in 1xPBS, and blocked in cold 4%BSA/PBS-1%Tween for 15 minutes. Sections were then incubated in 1% Wheat Germ Agglutin Rhodamine (RL_1022) in 4%BSA/PBS/1%Tween for 40 minutes in the dark at room temperature followed by additional 3×10 minute washes in PBS-1%Tween. Sections were mounted with Vectashield Antifade Mounting Medium with DAPI (Vector laboratories #93952-25). Images were gathered at 40x resolution with a Zeiss Axioplan 2e. 50-100 muscle fibers were randomly selected and manually segmented for cross-sectional area. Mean cross-sectional area from two technical replicates were used for statistical analysis.

### Myofibrillar extraction and electrophoresis

The myosin and actin fractionation methods were conducted, with slight modifications, as described in Roberts et al. (2020). In brief, 20mg of dorsal muscle tissue was dissected and immediately submerged in 500μl of homogenization buffer (98% RIPA Lysis Extraction Buffer (ThermoFisher #89900), 1% Halt Protease/Phosphatase Inhibitor Cocktail (Thermo Scientific #78442), 1% EDTA). The tissue was then homogenized with triple pure M-Bio grade high impact zirconium beads in a Beadbug 6 microtube homogenizer bead beater at 4900 RPM for 60 seconds followed by gentle agitation at 4°C for 60 minutes. The sample was then centrifuged at 4°C at 3,000 x g for 30 minutes. The supernatant (cytosolic fraction) was removed and frozen at −80°C. The pellet was resuspended with 500μl of homogenization buffer, centrifuged for 10 minutes at 3000 x g, and the supernatant was removed followed by two additional washes. The remaining myofibrillar pellet was then immediately frozen and placed at −80°C until further processing. The frozen myofibrillar pellet was resuspended in 125μl of homogenization buffer. 10μl of the resuspended pellet was added to 65μl of ddH_2_O and 25μl of 4x NuPage sample buffer, heated for 10 minutes at 70°C, and allowed to cool for 15 minutes at room temperature. Samples were loaded at equal volumes on a 10% SDS gel and ran at a current of 300 volts for 40 minutes at room temperature. Following electrophoresis, gels were washed in ddH_2_O for 10 minutes, incubated in Imperial Protein Stain (Thermo Scientific #24615) for 2 hours, and then washed in ddH2O for 90 minutes. The gels were digitally imaged with a Cannon EOS Rebel T7i Rebel with a Commander Optics Pro CPL 58mm Filter and pixel intensities of myosin and actin bands were quantified via ImageJ. Mouse skeletal muscle was processed identical to that described above to serve as an internal positive control for actin and myosin weight/band position.

### Muscle triglyceride quantification

Overnight fasted surface fish and Pachón cavefish were euthanized via ms222 and skeletal muscle immediately posterior to the dorsal fin was dissected. All muscle samples were immediately processed as described in the commercial triglyceride assay kit (#65336 - Sigma) on a 200 Pro Microplate Reader (TECAN). Due to extremely high triglyceride levels in all Pachón samples, repeated dilutions were needed to remain within the working range. However, three Pachón cavefish remained out of range (high triglyceride content) and were thus removed from triglyceride quantification. This did not interfere with the final interpretation of triglyceride levels between surface fish and Pachón cavefish.

### Muscle glycogen quantification

Skeletal muscle from the same fish used for triglyceride quantification were used to measure glycogen content following the protocol within the Glycogen Assay Kit (ab65620) on a 200 Pro Microplate Reader (TECAN). Similar to triglyceride quantification, a Pachón cavefish remained out of range following repeated dilutions resulting in this sample being discarded. This did not interfere with final interpretation of glycogen levels between surface fish and Pachón cavefish.

### Fish buoyancy

Fish buoyancy was conducted as described in Eastman (2020). In brief, adult fish were euthanized in ms222 and patted dry. All fish had their swim bladder removed of air and female fish had their eggs removed. Fish were then suspended from a hook that was attached to a weighing scale and their dry weight was measured. Fish were then submerged in water with all air removed and their weight was measured. The weight within the water was taken relative to their dry weight resulting in their percent water weight.

### Lab fish RNA-sequencing

All fish used were adult and housed at similar tank densities. Fish were fasted overnight and collected at 09:00 the following morning. Fish were euthanized in 500mg/L of ms222. Skeletal muscle was extracted immediately posterior to the dorsal fin and snap frozen in liquid nitrogen. ~100mg of frozen tissue was homogenized in 1mL Trizol (Ambion) with triple pure M-Bio grade high impact zirconium beads in a Beadbug 6 microtube bead beater. RNA was then extracted using standard phenol/chloroform extraction. The subsequent RNA pellet was cleaned with the RNeasy Mini Kit (Qiagen #74104) with on-column DNAse digestion (Qiagen #79256). Libraries were prepared according to the manufacturer’s instructions using the TruSeq Stranded mRNA Prep Kit (Illumina #20020594). The resulting libraries were quantified using a Bioanalyzer (Agilent Technologies) and Qubit fluorometer (Life Technologies). Libraries were normalized, pooled, multiplexed, and sequenced on an Illumina NextSeq 500 instrument as v2 Chemistry High Output 75bp single read runs. Following sequencing, raw reads were demultiplexed into Fastq format allowing up to one mismatch using Illumina bcl2fastq2 v2.18. Reads were aligned to UCSC genome astMex_2.0 with STAR aligner (version 2.7.3a) using Ensembl 102 gene models. TPM values were generated using RSEM (version v1.3.0). Pairwise differential expression analysis was performed using Bioconductor package edgeR (3.24.3 with R 3.5.2). Only protein coding genes and long non-coding RNAs (lncRNAs) were considered from the Ens_102 annotation. Only genes with counts per million expression ≥2 in at least 2 samples were kept for further analysis. Statistical significance was determined by fold change cutoff of 2 and false discovery rate (FDR) cutoff of 0.01 or 0.05. Principal Component Analysis (PCA) was performed using R package ‘stats’ using the prcomp function with the first two principal components shown. Gene Ontology (GO Term) enrichment was completed using TERMS2GO, an in-house R Shiny app (versions R 4.1.0, shiny 1.7.1). Significant gene ontology terms were identified using clusterProfiler’s enrichGO function (version 4.0.0) with AnnotationHub’s species database (version 3.0.0). GO Terms with adjusted p-values less than 0.01 and 0.05 were considered significant. Figures were generated using ggplot2 (version 3.3.5) and plotly (4.10.0).

### RT-qPCR

Following an overnight fast, skeletal muscle was extracted from *A. mexicanus* and immediately snap frozen. ~100mg of frozen tissue was homogenized in 1mL Trizol (Ambion) with triple pure M-Bio grade high impact zirconium beads in a Beadbug 6 microtube bead beater. RNA was then extracted using standard phenol/chloroform extraction. The subsequent RNA pellet was cleaned with the RNeasy Mini Kit (Qiagen #74104) with on-column DNAse digestion (Qiagen #79256). 1μg of quality RNA was then converted to cDNA with the iScript cDNA synthesis kit (Bio-Rad #1708890). 1-5ng of cDNA was used for qPCR utilizing SYBR green technology (Quantabio PerfeCTa SYBR Green FastMix Low Rox #66188573) with a QuantStudio 5 Real-Time PCR System. Specificity of each amplicon was confirmed via analysis of post-reaction dissociation curves, validating a single amplicon for each set of primers. Analysis was conducted using the Delta Delta *C_t_* method. All samples were run in triplicate and normalized to the housekeeping gene *rpl13a.* Primer sequences used are as follows:

*fast-myhc; ENSAMXG00000038006:*
FW: 5’ - TTCTTCTTGCCTCCCTTGCC-3’
RV: 5’-AAGGCTGAAGCCCACTTCTT -3’
*rpl13a; ENSAMXG00005024453:*
FW 5’-5’-GTTGGCATCAACGGATTTGG -3’
RV: 5’ - CCAGGTCAATGAAGGGGTCA -3’
*pparγ; ENSAMXG00000043041:*
FW: *5*’-GTCACCGCGATTCCTCTGAT-3’
RV: *5*’-ATCCCATGGGCCAGGAAAAC-3’.

### Muscle ex vivo kinetics

Muscle mechanics experiments were carried out at Widener University. For muscle mechanics experiments, fish were sacrificed and hypaxial myotomal muscle was dissected from between the pectoral and pelvic girdles. The extracted muscle was maintained in physiological saline (Seebacher et al. 2012) at room temperature. Muscle bundles consisted of 2-3 myomeres with a typical live muscle length of 4-5 mm and live muscle cross-sectional area of ~1 mm^2^. Isolated live bundles were tied with silk thread into a muscle mechanics chamber with a servomotor at one end (Aurora Scientific 318B) and force transducer at the other (Aurora Scientific 404A). The temperature-controlled chamber had platinum electrodes along each side. Experimental control was carried out using the Dynamic Muscle Control and Analysis Software (Aurora Scientific 615A) at 25°C. Sample size for muscle mechanics was n = 8 for groups of fish. Up to two muscle bundles could be examined per fish. Muscle length and stimulation conditions were optimized for each bundle to produce maximum tetanic force. Muscle bundles generating low force (<10 mN mm^-2^), indicating low tissue quality, were eliminated from the dataset. Isometric twitch and tetanic contractions were recorded and analyzed for time to maximum force (time from stimulus until maximum isometric force) and for ½ relax time (time from maximum force to 50% relaxation). Isovelocity experiments were used to measure the force-velocity relationship to determine muscle maximum shortening velocity (V_max_). A series of isovelocity ramps of increasing velocity were imposed on muscle bundles, and muscle tension was recorded during each ramp to plot the force-velocity curve for each bundle. Correcting for passive tension, a V_max_ was calculated by fitting the Hill muscle model (Seow, 2013) using Igor Pro (WaveMetrics). Following physiological measurements, bundles were stained in Trypan Blue (for dead tissue) in saline for 30 minutes at room temperature, embedded in gelatin (15% gelatin in saline) and frozen in dry-ice chilled hexanes. Live muscle area was then determined as described previously (Coughlin, 2000).

### Incremental swimming test

The incremental swim test was performed at the University of North Texas within a Blazka-style 1500mL Loligo Systems swim tunnel (#SW10040). Fish in each group were placed in the tunnel for a 20 minute acclimatization period with the water flow set at 5cm/s. This did not cause any strenuous movement or behavior from any of the fish. Fish within the “control” group were immediately removed following the 20 minute acclimatization period and euthanized. All other fish immediately followed the acclimatization period with the incremental swimming test consisting of ramping the water flow 5cm/s every 5 minutes. If the entire 5 minutes at a given water flow was completed, the flow would then increase in velocity an additional 5cm/s. The water flow was immediately turned off when the fish reached fatigue. Fatigue was defined as the fish remaining at the rear of the swim tunnel for 5 seconds, no longer swimming. Critical swimming speed (Ucrit) was measured with the following equation: [Uf + (T/t)dU]/cm, where Uf (cm s-1) is the highest swim velocity maintained for a full interval, T (s) is the time spent at the final velocity, t is the time interval (s), and dU is the increment in swim speed (cm s-1). Acute post fish were removed and immediately euthanized in ms222. 1-hour post fish were removed and placed in a 3L water tank for 1 hour. Following the hour, fish were euthanized in ms222. The incremental swim test protocol was approved by the UNT IACUC #19-011.

### Glucose measurements

Exercised and non-exercised fish were euthanized via ms222 followed by the caudal fin being removed and blood drawn directly to the AlphaTRAK Blood Glucose Monitoring System.

### Periodic acid-Schiff stain

Exercised and non-exercised fish had their skeletal muscle dissected immediately anterior to that obtained for the phosphoproteomic analysis and placed in 4% PFA at room temperature for 24 hours. Samples were washed 3×15 minutes in 1xPBS followed be serial dehydration in ethanol-1xPBS and placed at 4°C for 96 hours. Tissue was paraffin embedded, sectioned, deparaffinized, and dehydrated to ddH_2_O. Slides were placed in 5% periodic acid for 10 minutes, rinsed in distilled water for 5 minutes, and placed in Schiff Reagent for 15 minutes at room temperature. Slides were then washed in warm water for 10 minutes. Sections were mounted in 50% glycerol/1xPBS and imaged on a VS120 virtual slide microscope (Olympus). Areas for further analysis were randomly chosen and vetted for their morphological integrity. Muscle fibers showing adequate morphology were manually demarcated and pixel intensity was measured in ImageJ. Two technical replicates were analyzed for the majority of biological samples unless tissue morphology was inadequate for accurate fiber tracing. The mean pixel intensity of the two technical replicates was then used for statistical analysis.

### pAMPK quantification

~30mg of frozen skeletal muscle was submerged in ice cold RIPA Lysis Extraction Buffer (ThermoFisher #89900) supplemented with 1% Halt Protease/Phosphatase Inhibitor Cocktail (Thermo Scientific #78442) and homogenized with triple pure M-Bio grade high impact zirconium beads in a Beadbug 6 microtube homogenizer. Muscle lysate was then gently agitated at 4°C for 2 hours and spun down at 16,000 x g for 20 minutes. The subsequent supernatant was transferred to a new tube and snap frozen in liquid nitrogen. Protein quantification was determined with a Pierce BCA Protein Assay Kit (Thermo #23227). Samples were combined with NuPage LDS sample buffer 4x (#2083421) and 5% 2-Mercaptoethanol (Sigma Aldrich #M6250) and heated for 10 minutes at 70°C. Samples were then cooled for an additional 10 minutes and 60μg of protein was loaded onto a 10% polyacrylamide gel and electrophoretically separated at 140 volts for 65 minutes at room temperature. The gel was then transferred to a PVDF membrane for 60 minutes at 235 mAMPS at 4°C. The membrane was then blocked in LI-COR blocking buffer and incubated in 1:500 mAb pAMPK (T172) (CST #40H9) antibody overnight at 4°C. Following overnight incubation, the membrane was washed 1×10min/2×5min in TBS-1%Tween followed by incubation in the dark with Goat anti-Rabbit IgG Secondary Antibody (1:5000) (LI-COR #AB_2721181) for 1 hour at room temperature. The membrane was then washed 3 additional times in TBS-1%Tween. The membrane was scanned with the Odyssey LI-COR Scanning system (Odyssey CLx). For total protein quantification, the membrane was incubated in Imperial Protein Stain (Thermo Scientific #24615) for 2 hours and washed in ddH2O for 1 hour followed by imaging with a Cannon EOS Rebel T7i Rebel with a Commander Optics Pro CPL 58mm Filter. Pixel intensity was gathered with ImageJ and analyzed for both pAMPK and total protein. Statistical analysis was conducted against the normalized pAMPK to total protein concentration.

### Phosphoglucomutase activity assay and CIP incubation

Skeletal muscle from overnight fasted fish was extracted posterior to the dorsal fin and immediately flash frozen in liquid nitrogen. Frozen skeletal muscle was homogenized in the provided buffer with the Phosphoglucomutase activity assay kit (ab155896 abcam) supplemented with protease and phosphatase inhibitors (CIP treated samples contained only protease inhibitors). Samples were then gently agitated at 4°C for 30 minutes, spun down, and protein measured using the Pierce BCA Protein Assay Kit (product# 23227). For CIP incubation, 10μg of protein from each sample was incubated in 100μl 1xCIP buffer and 2μl CIP (5,000 U/mL) (M0525S NEG) for 30 minutes at 37°C. Phosphoglucomutase activity of CIP-treated and non-treated samples were then measured according to the manufacturer’s instructions (ab155896 abcam) using a 200 Pro Microplate Reader (TECAN).

### Transmission electronic microscopy (TEM)

For TEM, surface and cavefish muscle tissues were dissected and fixed with 50 mM Sodium Cacodylate (pH 7.4) containing 2.5% Paraformaldehyde and 2% Glutaraldehyde. The tissue blocks were post fixed with 2% OsO4 for 2 hours, and 1% Uranyl Acetate overnight. After dehydration with a graded Ethanol series, samples were infiltrated and embedded into Epon resin (EMS, Fort Washington, PA). Ultrathin (80 nm) sections were collected on copper grids, stained with 4% Uranyl Acetate in 75% Ethanol and 2% Lead Citrate. Sections were imaged using a FEI transmission electron microscope at 80kV. For image quantification, myofibrils were first identified in TEM images using un-trained Cellpose (Stringer et al. 2021). A subset of the results were hand corrected in Fiji using ROI tools, and these were used as training data for a standard Unet using DeepFiji (Nuckolls et al. 2020). This Unet model then performed the final segmentation of myofibril regions of ~125 images per sample. Data aggregation and plotting were done in python.

#### Global/Phosphorylated peptide extraction

##### Protein digestion

Skeletal muscle tissue from each timepoint were immediately dissected following euthanization via submersion in ms222 (500mg/L), frozen in liquid nitrogen, and stored at −80°C until further processing. For tissue homogenization and protein extraction, 30mg of frozen skeletal muscle was submerged in 1mL ice-cold lysis buffer containing 100mM triethylammonium bicarbonate (TEAB) and 10% SDS with 1x protease/phosphatase inhibitor cocktail and homogenized with triple pure M-Bio grade high impact zirconium beads in a Beadbug 6 microtube homogenizer. The lysate was then centrifuged at 16,000 x g for 10 minutes at 4°C. The supernatant was transferred to a new tube and measured for protein concentration with the Pierce BCA Protein Assay Kit (Thermo #23227). Following the recommended protocol for sample preparation before tandem mass tag (TMT, Thermo) labeling, 100μg of protein was transferred to a new tube and adjusted to a final volume of 100μl with 100mM TEAB. 5μl of 200mM tris(2-carboxyethyl)phosphine (TCEP) (Thermo Scientific #PI20490) was added and incubated at 55°C for 1 hour. 5μl of 500mM 2-chloroacetamide was then added to the sample and incubated for 30 minutes, protected from light at room temperature. Proteins were then precipitated by methanol/chloroform extraction. Protein pellets were resuspended in 40μl of 100mM TEAB and digested overnight at 37°C with shaking by the addition of 2μl of 1μg/μl trypsin (Promega, V5111) and 1μl of 80mM CAM (2mM final) (Sigma, C0267). Peptides were quantitated by the Pierce quantitative fluorometric peptide assay and stored at −20°C until labeling.

##### Tandem Mass Tag labeling

Prior to TMT labeling, each sample was spiked with a digest of pig serum albumin as an internal control. TMT 6-plex reagents were resuspended according to the protocol and 200μg was added to each sample. Samples were labeled for 1 hour at room temperature. To check labeling efficiency, 2μl of each reaction was diluted to 25μl with buffer A (5% acetonitrile (ACN), 0.1% formic acid (FA)) and injected via the autosampler of a Dionex Ultimate 3000 RSCL nano-HPLC onto a trapping column (Acclaim PepMap 100, 5μm particle size, 0.3mm x 5mm) using the loading pump flowing at 2.5 μl /min. Trapped peptides were eluted onto an in-house pulled separating column (1.9 μm resin (ReproSil, Dr. Maisch), 20cm packed into 75μm I.D. x 360μm O.D. fused silica) at 42°C with a flow rate of 250nL/min directly interfaced with an Thermo Orbitrap Eclipse mass spectrometer equipped with a FAIMS source at 2.5kV spray voltage. Peptides were loaded with 5% buffer B (80% ACN, 0.1%FA) and washed for 30 minutes before being eluted by a linear gradient to 40% B over 1 hour gradient. To quench the TMT reaction, 1μl of 5% hydroxylamine (Sigma) was added to each sample. The samples were combined to make four 6-plex technical replicates. Approximately 5% of the reaction was reserved for global TMT analysis and the remaining 95% was used for phospho-peptide enrichment. Phospho-peptides were enriched using Sequential enrichment from Metal Oxide Affinity Chromatography (SMOAC) (Jae Choi et al. 2017) approach in which phospho-peptides are sequentially enriched by TiO2 (High-Select TiO2 Phospho-peptide Enrichment Kit, Thermo) followed by FeNTA (High-Select Fe-NTA Phospho-peptide Enrichment Kit, Thermo). Peptides enriched for phospho-peptides were dried in a speed vacuum concentrator and stored at −20°C until resuspension and analysis.

##### Mass spectrometry analysis

Dried peptide samples were resuspended in 10μl or 25μl buffer A for global and phospho-enriched analyses, respectively. The unenriched global samples were further diluted 25x in buffer A before analysis. Samples were analyzed by LC/MSMS on an Eclipse mass spectrometer (Thermo Scientific) equipped with FAIMS and coupled to a Dionex Ultimate 3000 RSCL nano-HPLC. For each analysis, 20μl of resuspended 6-plex sample was injected onto a trapping column (Acclaim PepMap 100, 5μm particle size, 0.3mm x 5mm) using the loading pump flowing at 2.5μl/min. Trapped peptides were eluted onto an in-house pulled separating column (1.9 μm resin (ReprosSil, Dr. Maisch), 75μm I.D. x 20cm) at 42°C with a flow rate of 250nL/min directly interfaced with the Eclipse mass spectrometer equipped with a FAIMS source at 2.5kV spray voltage. Peptides were loaded with 5% buffer B (80% ACN, 0.1%FA) and washed for 30 minutes before a linear gradient to 40% B over 3 hour or 4 hour for phospho-enriched or global samples, respectively. The gradient was increased to 85% B over 10 minutes and held for an additional 5 minutes before re-equilibration for 30 minutes at 5% buffer B for the next injection. Peptides were analyzed using the TMT-SPS-MS3 with FAIMS method on the Eclipse. Briefly, peptides were scanned in the Orbitrap at 120,000 resolving power before MS2 fragmentation by CID at 35% NCE and detection in the ion trap set to rapid detection. Synchronous precursor scanning (SPS) selected the top 10 MS2 peptides for TMT reporter ion detection in the Orbitrap using HCD fragmentation at 65% NCE at 50,000 resolving power. Top speed of 1s for each FAIMS compensation voltage (CV) at −40V, −60V, and −80V was used. Dynamic exclusion was enabled for 45s to deepen the proteome coverage.

##### Database searching

RAW files were directly uploaded to Proteome Discoverer v. 2.4. Samples were searched against the *Astyanax mexicanus* database (downloaded from NCBI 04-25-2019) and a database containing 426 common contaminants with static peptide N-terminal TMT reporter ion modification (+229.163 Da) and cysteine alkylation (+57.125 Da) and variable modification search for methionine oxidation (+15.995 Da), serine, threonine, and tyrosine phosphorylation (+79.993 Da) and lysine TMT modification (+229.163 Da). Peptides were normalized to total peptide amount for the global analysis. In the phospho-peptide enriched samples, only peptides containing phosphorylation were quantitated. False discovery rates for the global dataset were maintained at 0.5%, 1%, and 6.8% for peptide spectrum matches, peptides, and proteins, respectively. False discovery rates for the phospho-enriched dataset were maintained at 0.3, 1%, and 4.1% for peptide spectrum matches, peptides, and proteins, respectively. 640 proteins were identified in the global dataset containing TMT reporter ions and identified in at least 2 replicates (Data S7). In the phospho-enriched dataset, there were 124 proteins and 430 phosphorylated peptides identified that were quantitated by TMT reporter ions in at least 2 replicates (Data S10). Mass spectrometry data were processed with MaxQuant (v1.5.2.10) and searched with Andromeda against either the mouse or rat UniProt database. KEGG Pathway Analysis was conducted in g.profiler (Raudvere et al. 2019) using the functional profiling g:GOSt pipeline against the Astyanax_mexicanus_2.0 Ensemble genome.

##### Targeted proteomics

An additional 10% of the remaining global protein samples were used for targeted proteomics. Samples were diluted 1μl to 25μl of Buffer A and was transferred to an autosampler vial. The targeted proteomics samples were analyzed with a similar TMT-SPS-MS3 method to the global analysis with the exceptions that the gradient was shortened to 3 hour and an inclusion list was used with 2-3 peptides from each of the 22 targeted proteins (Data S9B). Dynamic exclusion was not used. Data were analyzed with Proteome Discoverer 2.4 with SEQUEST-HT against the *Astyanax mexicanus* protein database as used in the global analysis. Data was searched to include static modifications of cysteine alkylation (+57.125 Da) and peptide N-terminal TMT tag (+229.163Da) and dynamic modification of lysine TMT modification (+229.163 Da), methionine oxidation (+15.995 Da), and serine, threonine, and tyrosine phosphorylation (+79.993 Da). Data was normalized in PD to the highest total TMT reporter abundance. It was observed that all samples in runs 1 through 3 clustered well within populations (i.e. cave and surface clustered separately) whereas both cavefish and surface samples in run 4 clustering together, independent of population. We reasoned this reflected poor running of the samples and as such samples in run 4 were removed and all subsequent analysis included only samples 1 through 3.

##### Data normalization for Principal Component Analysis

For each biological replicate, phospho-proteomic data was quantified as mean peak intensity of each modified peptide. Peptide level quantities exhibited severe biases and dropouts, with the strength of these effects depending on the biological replicate of the measurement. In order to visualize biological variation, rather than batch-related effects, we attempted to correct for both dropouts and biases. In order to correct for dropouts, we used hard imputation from the *filling* R package with the default rank of min(dim(A))-1, which evaluates to 23 for phospho-peptide datasets. Using this imputed data, we performed batch-normalization using the Removal of Unwanted Variation (RUV) R package (Jacob et al. 2016). Briefly, we used the RUVIII function with a k-factor of 10. RUV makes use of so-called “negative controls,” i.e. features that are not expected to change across experiments. Due to the comparatively understudied nature of our *A. mexicanus* model system, we chose to model all peptides as negative controls, a conservative approach often used when candidate negative controls are not available. Finally, these imputed and normalized values were used to compute a PCA decomposition based on standard routines in the Python *scikit-learn* package.

##### Statistical modeling and ranking

In order to determine phosphorylation-related changes associated with the cave adaptation phenotype, we sought to identify peptides with significant differences in either 1) differential phosphorylation as a function of physiological state (pre vs acute swimming state or pre vs 1-hour post-challenge state), or 2) baseline measurements for all three physiological timepoints. We further sought to identify peptides with a large differential or baseline change in one population (surface or cave) but not the other, thereby suggesting a potential fitness advantage of the peptide in the cave or surface environment, but not both. In order to accomplish this, we did not use the normalized data as described above, but rather used a linear mixed-effect model to estimate the contribution due to batch effects. Briefly, we constructed a model incorporating peptide / physiological timepoint interaction as a fixed effect and mean peak intensity per replicate as a random effect (for measuring baseline differences between populations, we instead used peptide/population interaction as the fixed effect). Fitting this model allowed us to compute statistical metrics for the differential abundance of a given phospho-peptide without confounding due to batch effects. We then ranked peptides from greatest to least differential abundance in Pachón for each timepoint and least to greatest abundance in surface (i.e. the ranking criteria for surface is reversed). We then merged these rankings using rank products according to (Breitling et al. 2004) to yield a ranking of peptides with a large timepoint differential in one population (Pachón) but not the other (surface). We repeated this process interchanging Pachón and surface to generate complementary rankings. Finally, we used a similar rank product approach to rank baseline-differences.

#### Data availability

The mass spectrometry proteomics data have been deposited to the ProteomeXchange Consortium (Perez-Riverol et al. 2019; Deutsch et al. 2020) via the PRIDE partner repository with the dataset identifier <u>PXD024165</u> and may be accessed through the MassIVE partner repository via ftp with username <u>MSV000086857</u> and password “OlsenMuscle2021”. The RNA-seq datasets can be found at the GEO accession number GSE196531. Original data underlying this manuscript may be accessed after publication from the Stowers Original Data Repository at http://www.stowers.org/research/publications/libpb-1679.

---

**Figure S1.**
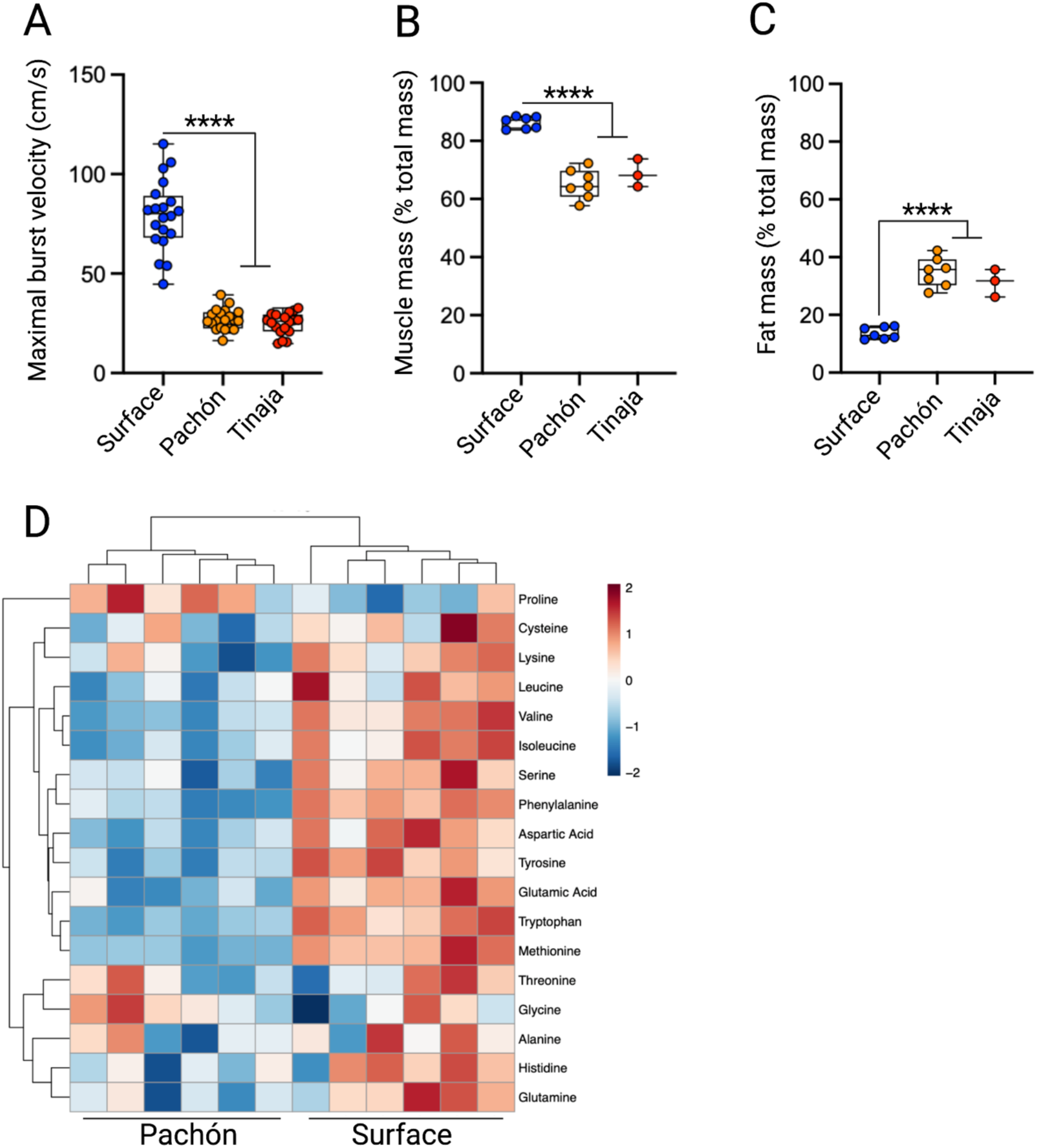
Maximal burst velocity and body composition of *A. mexicanus*. (A) Maximal burst velocity of the surface fish (n=20) and two independently evolved cavefish populations: Pachón (n=20) and Tinaja (=18). Relative (B) muscle mass and (C) fat mass of histological sections from surface fish (n=7), Pachón (n=7), and Tinaja (n=3) cavefish following echoMRI analysis. (D) Heatmap of the identified amino acids (18 in total) within Pachón cavefish and surface fish. Arginine and Asparagine were not detected. Data was used from the following shiny app: https://cavefin.shinyapps.io/shiny (Medley et al. 2020). Significance was calculated with a one-way ANOVA with Bonferroni correction for Fig. S1A-C. Data is presented as ±SEM, \*\*\*\**p*<0.0001.

**Figure S2.**
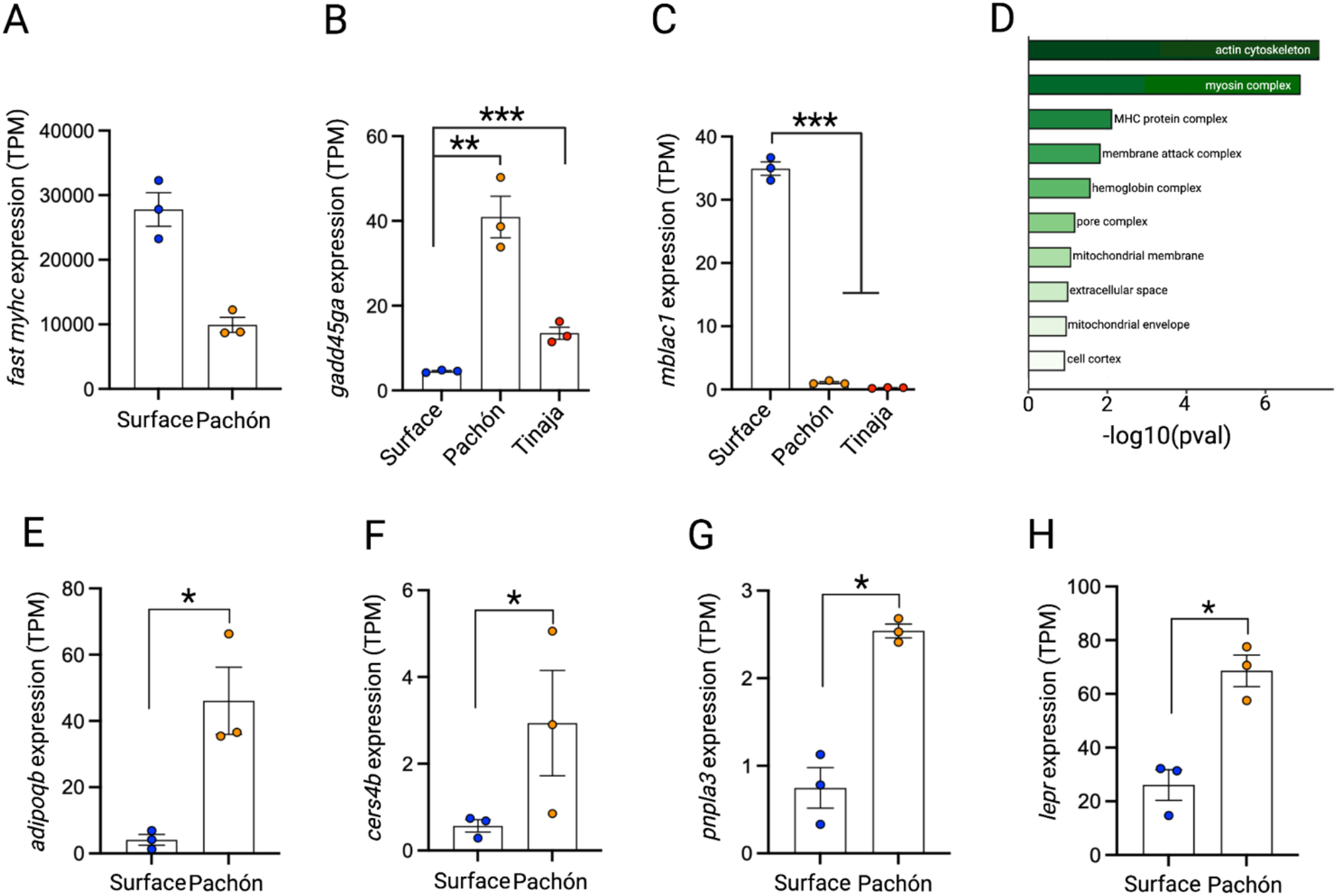
Gene expression of the *A. mexicanus* muscle tissue. (A) Cumulative gene expression in transcripts per million (TPM) of all fast myosin heavy chain (*fast-myhc*) transcripts significantly different between surface fish and Pachón cavefish. Gene expression in TPM of (B) *gadd45ga* and (C) *mblac1* (n=3 per population). (D) GO-enrichment analysis of the DEG’s conserved in both wild-caught and laboratory-reared *A. mexicanus*. Gene expression in TPM of (E) *adipoqb* (F) *cers4b*, (G) *pnpla3,* and (H) *lepr* (n=3 per population). Statistical analysis for RNA-seq can be found in the methods. Data is presented as ±SEM, \*\*\**p*<0.001.

**Figure S3.**
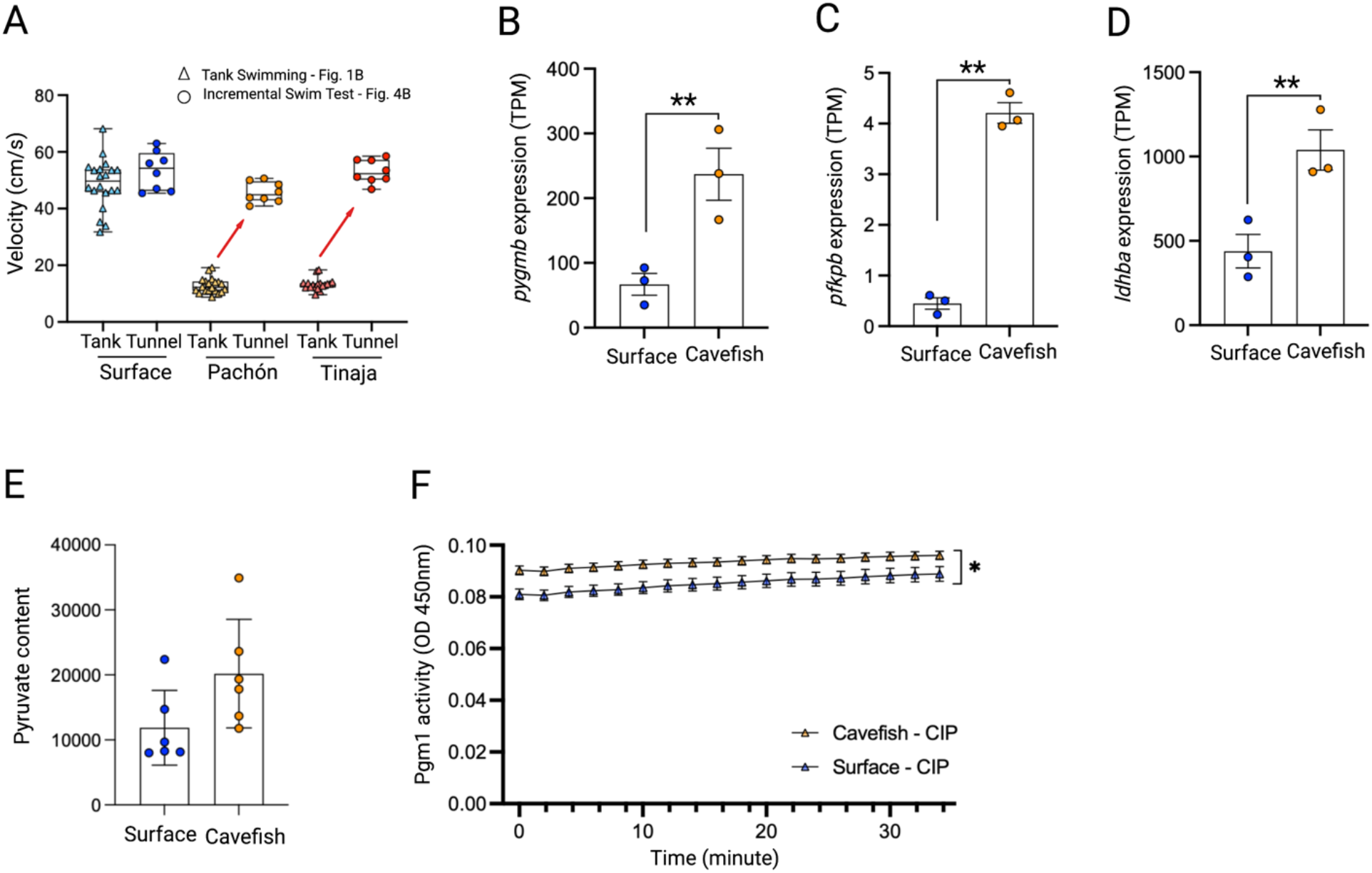
Swimming velocity and metabolic investment within the *A. mexicanus*. (A) Comparison of the maximal swimming velocity reached during the incremental swim test (circles) and average burst velocity as shown in Figure 1B (triangles). The red arrow highlight the change in swimming velocity within the cavefish (Pachón and Tinaja). Change in gene expression (TPM = transcript per million) of (B) *pygmb*, (C) *pfkpb*, and (D) *ldhba*. (E) Pyruvate content in the skeletal muscle of surface fish and cavefish (Pachón) (n=6 per population). (F) Pgm1 activity within surface fish and cavefish (Pachón) following CIP incubation (n=6 per population). While at baseline cavefish have increased Pgm1 fluorescence, their activity over time is less than that in surface fish. For Fig. S3H, significance was calculated with a two-way repeated measures ANOVA with Benjamini and Hochberg FDR correction. Statistical analysis for RNA-seq data can be found in the methods. Data is presented as ±SEM, \**p*<0.05, \*\**p*<0.01.

## References

Aspiras, A. C., Rohner, N., Martineau, B., Borowsky, R. L., & Tabin, C. J. (2015). Melanocortin 4 receptor mutations contribute to the adaptation of cavefish to nutrient-poor conditions. Proc Natl Acad Sci U S A, 112(31), 9668–9673. doi:10.1073/pnas.1510802112

Balázs, G., Lewarne, B., & Herczeg, G. (2020). Extreme site fidelity of the olm (Proteus anguinus) revealed by a long-term capture-mark-recapture study. Journal of Zoology, 311(2), 99–105. doi.org/10.1111/jzo.12760

Batista, T. M., Jayavelu, A. K., Wewer Albrechtsen, N. J., Iovino, S., Lebastchi, J., Pan, H.,… Kahn, C. R. (2020). A Cell-Autonomous Signature of Dysregulated Protein Phosphorylation Underlies Muscle Insulin Resistance in Type 2 Diabetes. Cell Metab, 32(5), 844–859 e845. doi:10.1016/j.cmet.2020.08.007

Biltz, N. K., Collins, K. H., Shen, K. C., Schwartz, K., Harris, C. A., & Meyer, G. A. (2020). Infiltration of intramuscular adipose tissue impairs skeletal muscle contraction. J Physiol, 598(13), 2669–2683. doi:10.1113/jp279595

Bodine, S. C. (2013). Disuse-induced muscle wasting. Int J Biochem Cell Biol, 45(10), 2200–2208. doi:10.1016/j.biocel.2013.06.011

Boggs, T., & Gross, J. (2021). Reduced Oxygen as an Environmental Pressure in the Evolution of the Blind Mexican Cavefish. Diversity, 13(1), 26.

Booth, F. W., Roberts, C. K., Thyfault, J. P., Ruegsegger, G. N., & Toedebusch, R. G. (2017). Role of Inactivity in Chronic Diseases: Evolutionary Insight and Pathophysiological Mechanisms. Physiol Rev, 97(4), 1351–1402. doi:10.1152/physrev.00019.2016

Breitling, R., Armengaud, P., Amtmann, A., & Herzyk, P. (2004). Rank products: a simple, yet powerful, new method to detect differentially regulated genes in replicated microarray experiments. FEBS Lett, 573(1-3), 83–92. doi:10.1016/j.febslet.2004.07.055

Carlson, B. M., Klingler, I. B., Meyer, B. J., & Gross, J. B. (2018). Genetic analysis reveals candidate genes for activity QTL in the blind Mexican tetra, *Astyanax mexicanus*. PeerJ, 6, e5189. https://doi.org/10.7717/peerj.518

Chen, L., Bai, Y., Everaert, N., Li, X., Tian, G., Hou, C., & Zhang, D. (2019). Effects of protein phosphorylation on glycolysis through the regulation of enzyme activity in ovine muscle. Food Chem, 293, 537–544. doi:10.1016/j.foodchem.2019.05.011

Choi, S. J., Files, D. C., Zhang, T., Wang, Z. M., Messi, M. L., Gregory, H.,… Delbono, O. (2016). Intramyocellular Lipid and Impaired Myofiber Contraction in Normal Weight and Obese Older Adults. J Gerontol A Biol Sci Med Sci, 71(4), 557–564. doi:10.1093/gerona/glv169

Contrepois, K., Wu, S., Moneghetti, K. J., Hornburg, D., Ahadi, S., Tsai, M. S.,… Snyder, M. P. (2020). Molecular Choreography of Acute Exercise. Cell, 181(5), 1112–1130 e1116. doi:10.1016/j.cell.2020.04.043

Coughlin, D. J. (2000). Power production during steady swimming in largemouth bass and rainbow trout. J Exp Biol, 203(Pt 3), 617–629. Retrieved from https://www.ncbi.nlm.nih.gov/pubmed/10637190

Deutsch, E. W., Bandeira, N., Sharma, V., Perez-Riverol, Y., Carver, J. J., Kundu, D. J.,… Vizcaino, J. A. (2020). The ProteomeXchange consortium in 2020: enabling ‘big data’ approaches in proteomics. Nucleic Acids Res, 48(D1), D1145–D1152. doi:10.1093/nar/gkz984

Dickinson, M. H., Farley, C. T., Full, R. J., Koehl, M. A., Kram, R., & Lehman, S. (2000). How animals move: an integrative view. Science, 288(5463), 100–106. doi:10.1126/science.288.5463.100

Dumesic, P. A., Egan, D. F., Gut, P., Tran, M. T., Parisi, A., Chatterjee, N.,… Spiegelman, B. M. (2019). An Evolutionarily Conserved uORF Regulates PGC1alpha and Oxidative Metabolism in Mice, Flies, and Bluefin Tuna. Cell Metab, 30(1), 190–200 e196. doi:10.1016/j.cmet.2019.04.013

Eastman, J. T. (2020). The buoyancy-based biotope axis of the evolutionary radiation of Antarctic cryonotothenioid fishes. Polar Biology, 43, 1217–1231.

Elipot, Y., Hinaux, H., Callebert, J., & Rétaux, S. (2013). Evolutionary shift from fighting to foraging in blind cavefish through changes in the serotonin network. Curr Biol, 23(1), 1–10. doi:10.1016/j.cub.2012.10.044

Fontes-Oliveira, C. C., Busquets, S., Fuster, G., Ametller, E., Figueras, M., Olivan, M.,… Argilés, J. M. (2014). A differential pattern of gene expression in skeletal muscle of tumor-bearing rats reveals dysregulation of excitation-contraction coupling together with additional muscle alterations. Muscle Nerve, 49(2), 233–248. doi:10.1002/mus.23893

Franchi, M. V., Reeves, N. D., & Narici, M. V. (2017). Skeletal Muscle Remodeling in Response to Eccentric vs. Concentric Loading: Morphological, Molecular, and Metabolic Adaptations. Frontiers in physiology, 8, 447. https://doi.org/10.3389/fphys.2017.00447

Fröbert, O., Frøbert, A. M., Kindberg, J., Arnemo, J. M., & Overgaard, M. T. (2020). The brown bear as a translational model for sedentary lifestyle-related diseases. J Intern Med, 287(3), 263–270. doi:10.1111/joim.12983

Gross, J. B. (2012). The complex origin of Astyanax cavefish. BMC Evol Biol, 12, 105. doi:10.1186/1471-2148-12-105

Hardaway, J. A., Sturgeon, S. M., Snarrenberg, C. L., Li, Z., Xu, X. Z., Bermingham, D. P.,… Blakely, R. D. (2015). Glial Expression of the Caenorhabditis elegans Gene swip-10 Supports Glutamate Dependent Control of Extrasynaptic Dopamine Signaling. J Neurosci, 35(25), 9409–9423. doi:10.1523/jneurosci.0800-15.2015

Hendrickson, D., Krejca, J., & Martínez, J. (2001). Mexican Blindcats Genus Prietella (Siluriformes: Ictaluridae): an overview of Recent Explorations. Environmental Biology of Fishes, 62, 315–337. doi:10.1023/A:1011808805094

Herman, A., Brandvain, Y., Weagley, J., Jeffery, W. R., Keene, A. C., Kono, T. J. Y.,… McGaugh, S. E. (2018). The role of gene flow in rapid and repeated evolution of cave-related traits in Mexican tetra, Astyanax mexicanus. Mol Ecol, 27(22), 4397–4416. doi:10.1111/mec.14877

Hermansen, L., Hultman, E., & Saltin, B. (1967). Muscle glycogen during prolonged severe exercise. Acta Physiol Scand, 71(2), 129–139. doi:10.1111/j.1748-1716.1967.tb03719.x

Hüppop, K. (2000). How do cave animals cope with food scarcity in caves? Amsterdam: Elsevier Press.

Jacob, L., Gagnon-Bartsch, J. A., & Speed, T. P. (2016). Correcting gene expression data when neither the unwanted variation nor the factor of interest are observed. Biostatistics, 17(1), 16–28. doi:10.1093/biostatistics/kxv026

Jae Choi, S. I. S., Ryan Bomgarden, John C. Rogers. (2017). Sequential enrichment from Metal Oxide Affinity Chromatography (SMOAC), a phosphoproteomicsstrategy for the separation of multiply phosphorylated from monophosphorylatedpeptides. ThermoFisher Scientific.

Kaji, T., Anker, A., Wirkner, C. S., & Palmer, A. R. (2018). Parallel Saltational Evolution of Ultrafast Movements in Snapping Shrimp Claws. Curr Biol, 28(1), 106–113 e104. doi:10.1016/j.cub.2017.11.044

Krishnan, J., Seidel, C. W., Zhang, N., VanCampen, J., Peuß, R., Xiong, S.,… Rohner, N. (2020). Genome-wide analysis of cis-regulatory changes in the metabolic adaptation of cavefish. bioRxiv, 2020.2008.2027.270371. doi:10.1101/2020.08.27.270371

Krishnan, J., Wang, Y., Kenzior, O., Huzaifa, H., Olsen, L., Tsuchiya, D.,… Rohner, N. (2022). Liver-derived cell lines from cavefish Astyanax mexicanus as an in vitro model for studying metabolic adaptation. bioRxiv, 2022.2001.2006.475101. doi:10.1101/2022.01.06.475101

Lee, Y., Stiers, K. M., Kain, B. N., & Beamer, L. J. (2014). Compromised catalysis and potential folding defects in in vitro studies of missense mutants associated with hereditary phosphoglucomutase 1 deficiency. J Biol Chem, 289(46), 32010–32019. doi:10.1074/jbc.M114.597914

Li, Y., Liang, R., Sun, M., Li, Z., Sheng, H., Wang, J.,… Shan, C. (2020). AMPK-dependent phosphorylation of HDAC8 triggers PGM1 expression to promote lung cancer cell survival under glucose starvation. Cancer Lett, 478, 82–92. doi:10.1016/j.canlet.2020.03.007

Maier, G., Delezie, J., Westermark, P. O., Santos, G., Ritz, D., & Handschin, C. (2021). Transcriptomic, proteomic and phosphoproteomic underpinnings of daily exercise performance and zeitgeber activity of training in mouse muscle. J Physiol. doi:10.1113/jp281535

Medley, JK., Persons, J., Peuss, R., Olsen, L., Xiong, S., Rohner, N., Untargeted Metabolomics of the cavefish Astyanax mexicanus Reveals the Basis of Metabolic Strategies in Adaptation to Extreme Conditions. Biorxiv. 2020.10.27.358077

Nuckolls, N. L., Mok, A. C., Lange, J. J., Yi, K., Kandola, T. S., Hunn, A. M.,… Zanders, S. E. (2020). The wtf4 meiotic driver utilizes controlled protein aggregation to generate selective cell death. Elife, 9. doi:10.7554/eLife.55694

Paz, A., McDole, B., Kowalko, J. E., Duboue, E. R., & Keene, A. C. (2020). Evolution of the acoustic startle response of Mexican cavefish. J Exp Zool B Mol Dev Evol, 334(7-8), 474–485. doi:10.1002/jez.b.22988

Perez-Riverol, Y., Csordas, A., Bai, J., Bernal-Llinares, M., Hewapathirana, S., Kundu, D. J.,… Vizcaino, J. A. (2019). The PRIDE database and related tools and resources in 2019: improving support for quantification data. Nucleic Acids Res, 47(D1), D442–D450. doi:10.1093/nar/gky1106

Raudvere, U., Kolberg, L., Kuzmin, I., Arak, T., Adler, P., Peterson, H., & Vilo, J. (2019). g:Profiler: a web server for functional enrichment analysis and conversions of gene lists (2019 update). Nucleic Acids Res, 47(W1), W191–W198. doi:10.1093/nar/gkz369

Riddle, M. R., Aspiras, A. C., Gaudenz, K., Peuss, R., Sung, J. Y., Martineau, B.,… Rohner, N. (2018). Insulin resistance in cavefish as an adaptation to a nutrient-limited environment. Nature, 555(7698), 647–651. doi:10.1038/nature26136

Rivas, D. A., Rice, N. P., Ezzyat, Y., McDonald, D. J., Cooper, B. E., & Fielding, R. A. (2019). Sphingosine-1-phosphate analog FTY720 reverses obesity but not age-induced anabolic resistance to muscle contraction. American Journal of Physiology-Cell Physiology, 317(3), C502–C512. doi:10.1152/ajpcell.00455.2018

Roberts, M. D., Haun, C. T., Vann, C. G., Osburn, S. C., & Young, K. C. (2020). Sarcoplasmic Hypertrophy in Skeletal Muscle: A Scientific “Unicorn” or Resistance Training Adaptation? Front Physiol, 11, 816. doi:10.3389/fphys.2020.00816

Roberts, M. D., Young, K. C., Fox, C. D., Vann, C. G., Roberson, P. A., Osburn, S. C.,… Kavazis, A. N. (2020). An optimized procedure for isolation of rodent and human skeletal muscle sarcoplasmic and myofibrillar proteins. J Biol Methods, 7(1), e127. doi:10.14440/jbm.2020.307

Rohner, N. (2018). Cavefish as an evolutionary mutant model system for human disease. Dev Biol, 441(2), 355–357. doi:10.1016/j.ydbio.2018.04.013

Rohner, N., Jarosz, D. F., Kowalko, J. E., Yoshizawa, M., Jeffery, W. R., Borowsky, R. L.,… Tabin, C. J. (2013). Cryptic variation in morphological evolution: HSP90 as a capacitor for loss of eyes in cavefish. Science, 342(6164), 1372–1375. doi:10.1126/science.1240276

Rome, L. C., Syme, D. A., Hollingworth, S., Lindstedt, S. L., & Baylor, S. M. (1996). The whistle and the rattle: the design of sound producing muscles. Proc Natl Acad Sci U S A, 93(15), 8095–8100. doi:10.1073/pnas.93.15.8095

Rudrappa, S. S., Wilkinson, D. J., Greenhaff, P. L., Smith, K., Idris, I., & Atherton, P. J. (2016). Human Skeletal Muscle Disuse Atrophy: Effects on Muscle Protein Synthesis, Breakdown, and Insulin Resistance-A Qualitative Review. Front Physiol, 7, 361. doi:10.3389/fphys.2016.00361

Salin, K., Voituron, Y., Mourin, J., & Hervant, F. (2010). Cave colonization without fasting capacities: an example with the fish Astyanax fasciatus mexicanus. Comparative biochemistry and physiology. Part A, Molecular & integrative physiology, 156(4), 451–457. https://doi.org/10.1016/j.cbpa.2010.03.030

Sänger, A. M., Stoiber, W. (2001). Muscle fiber diversity and plasticity. Fish Physiol., 18, 187–250.

Schiaffino, S., & Reggiani, C. (2011). Fiber types in mammalian skeletal muscles. Physiol Rev, 91(4), 1447–1531. doi:10.1152/physrev.00031.2010

Seebacher, F., Pollard, S. R., & James, R. S. (2012). How well do muscle biomechanics predict whole-animal locomotor performance? The role of Ca2+ handling. J Exp Biol, 215(Pt 11), 1847–1853. doi:10.1242/jeb.067918

Seow, C. Y. (2013). Hill’s equation of muscle performance and its hidden insight on molecular mechanisms. J Gen Physiol, 142(6), 561–573. doi:10.1085/jgp.201311107

Stojkovic, T., Vissing, J., Petit, F., Piraud, M., Orngreen, M. C., Andersen, G.,… Laforêt, P. (2009). Muscle glycogenosis due to phosphoglucomutase 1 deficiency. N Engl J Med, 361(4), 425–427. doi:10.1056/NEJMc0901158

Stringer, C., Wang, T., Michaelos, M., & Pachitariu, M. (2021). Cellpose: a generalist algorithm for cellular segmentation. Nat Methods, 18(1), 100–106. doi:10.1038/s41592-020-01018-x

Tegtmeyer, L. C., Rust, S., van Scherpenzeel, M., Ng, B. G., Losfeld, M. E., Timal, S.,… Marquardt, T. (2014). Multiple phenotypes in phosphoglucomutase 1 deficiency. N Engl J Med, 370(6), 533–542. doi:10.1056/NEJMoa1206605

van der Weele, C. M., & Jeffery, W. R. (2022). Cavefish cope with environmental hypoxia by developing more erythrocytes and overexpression of hypoxia inducible genes. Elife, 11. doi:10.7554/eLife.69109

Wisdom, K. M., Delp, S. L., & Kuhl, E. (2015). Use it or lose it: multiscale skeletal muscle adaptation to mechanical stimuli. Biomech Model Mechanobiol, 14(2), 195–215. doi:10.1007/s10237-014-0607-3

Xiong, S., Wang, W., Kenzior, A., Olsen, L., Krishnan, J., Persons, J.,… Rohner, N. (2021). Enhanced lipogenesis through Pparγ helps cavefish adapt to food scarcity. bioRxiv, 2021.2004.2027.441667. doi: 10.1101/2021.04.27.441667

